# A molecular portrait of microsatellite instability across multiple cancers

**DOI:** 10.1101/079152

**Authors:** Isidro Cortes-Ciriano, Sejoon Lee, Woong-Yang Park, Tae-Min Kim, Peter J. Park

## Abstract

Microsatellite instability (MSI) refers to the hypermutability of the cancer genome due to impaired DNA mismatch repair. Although MSI has been studied for decades, the large amount of sequencing data now available allows us to examine the molecular fingerprints of MSI in greater detail. Here, we analyze ~8000 exome and ~1000 whole-genome pairs across 23 cancer types. Our pan-cancer analysis reveals that the prevalence of MSI events is highly variable within and across tumor types including some in which MSI is not typically examined. We also identify genes in DNA repair and oncogenic pathways recurrently subject to MSI and uncover non-coding loci that frequently display MSI events. Finally, we propose an exome-based predictive model for the MSI phenotype that achieves high sensitivity and specificity. These results advance our understanding of the genomic drivers and consequences of MSI, and a comprehensive catalog of tumor-type specific MSI loci we have generated enables efficient panel-based MSI testing to identify patients who are likely to benefit from immunotherapy.

## Introduction

Microsatellites (MS) are tandem repeats of short DNA sequences, from mononucleotides to longer repeats such as hexanucleotides. MS are abundant throughout the genome and, owing to their high mutation rates, have been widely used as polymorphic genetic markers in population genomics and forensics. Microsatellite instability (MSI) is a hypermutator phenotype that occurs in tumors with impaired DNA mismatch repair (MMR) and is characterized by widespread length polymorphisms of MS repeats due to DNA polymerase slippage^1^ as well as by elevated frequency of single-nucleotide mutations (SNVs). MSI is caused by inactivation of MMR genes (e.g., *MLH1*, *MSH2*, *MSH3*, *MSH6* and *PMS2*) through somatic mutations in sporadic cases, with increased risk of cancer for those with inherited germline mutations (i.e., Lynch syndrome)^2^. MSI also occurs by hypermethylation of the *MLH1* promoter (e.g., associated with the somatic *BRAF* V600E mutation)^3^, epigenetic inactivation of *MSH2*^4^, or down-regulation of MMR genes by microRNAs^5^. MSI events within coding regions can alter the reading frame and lead to truncated, functionally-impaired proteins.^6^

MSI is observed in 15% of sporadic colorectal tumors diagnosed in the United States^7^, and has been reported in glioblastomas, lymphomas, stomach, urinary tract, ovarian and endometrial tumors^8^. In clinical settings, detection of MSI is customarily performed by immunohistochemical analysis of MMR proteins or by profiling the Bethesda markers^7^, which often include two mononucleotide (BAT25 and BAT26) and three dinucleotide (D5S346, D2S123 and D17S250) MS loci. Colorectal tumors unstable at >40% of the Bethesda markers are considered MSI-High (MSI-H) and are known to have better a prognosis and to be less prone to metastasis than MSI stable (MSS) tumors^9^.

It was conjectured more than two decades ago that the less aggressive nature of MSI tumors may be due to their high incidence of somatic mutations, which results in a greater likelihood of having mutated genes whose products elicit antitumor immune responses^10^. Indeed, in melanoma and lung tumors, an elevated mutational load has been associated with an increased rate of response to anti-CTLA-4 and anti-PD-1 therapies, respectively, likely as a result of a higher neo-antigen burden leading to antitumor immune response^11,12^. Other reports have shown that colorectal patients with MMR deficiency have better responses to immunotherapy by PD-1 immune checkpoint blockade and show improved progression-free survival^13^. Although the precise link between the mutator phenotype with MSI and patient response to immunotherapy remains to be elucidated, it is clear that accurate identification of patients with the hypermutator phenotype and their genomic characterization is of therapeutic importance.

In this study, we utilize data from The Cancer Genome Atlas (TCGA)^14^ to expand our previous analysis of MSI in 277 colorectal and uterine endometrial exomes to a pan-cancer cohort comprising ~8,000 exomes and ~1000 whole genomes, spanning 23 tumor types. We systematically profile the patterns of MSI mutations in both nuclear and mitochondrial DNA, characterize the affected pathways, and find association with epigenomic features. These analyses uncover new genes harboring frameshift MSI events with varying degrees of cancer-type specificity and generate the most comprehensive catalogue of MS loci selectively subject to DNA slippage events in MSI-H tumors to date. This set includes loci in the non-coding portions of the genome revealed by whole-genome sequencing. Lastly, we describe highly accurate predictive models of MSI-H status based on exome sequencing data.

## Results

### The exome-wide profiles of MSI in cancer genomes

To obtain an MSI landscape in cancer patients, we analyzed exome sequencing data for 7,919 tumor and matched normal pairs across 23 cancer types (Table 1). We identified 386,396 microsatellite repeats among the 39,496 RefSeq mRNA sequences^15^ and tested for the presence of MSI at microsatellites that had sufficient coverage in the exome data (Fig. 1a; Online Methods).

**Table 1.**
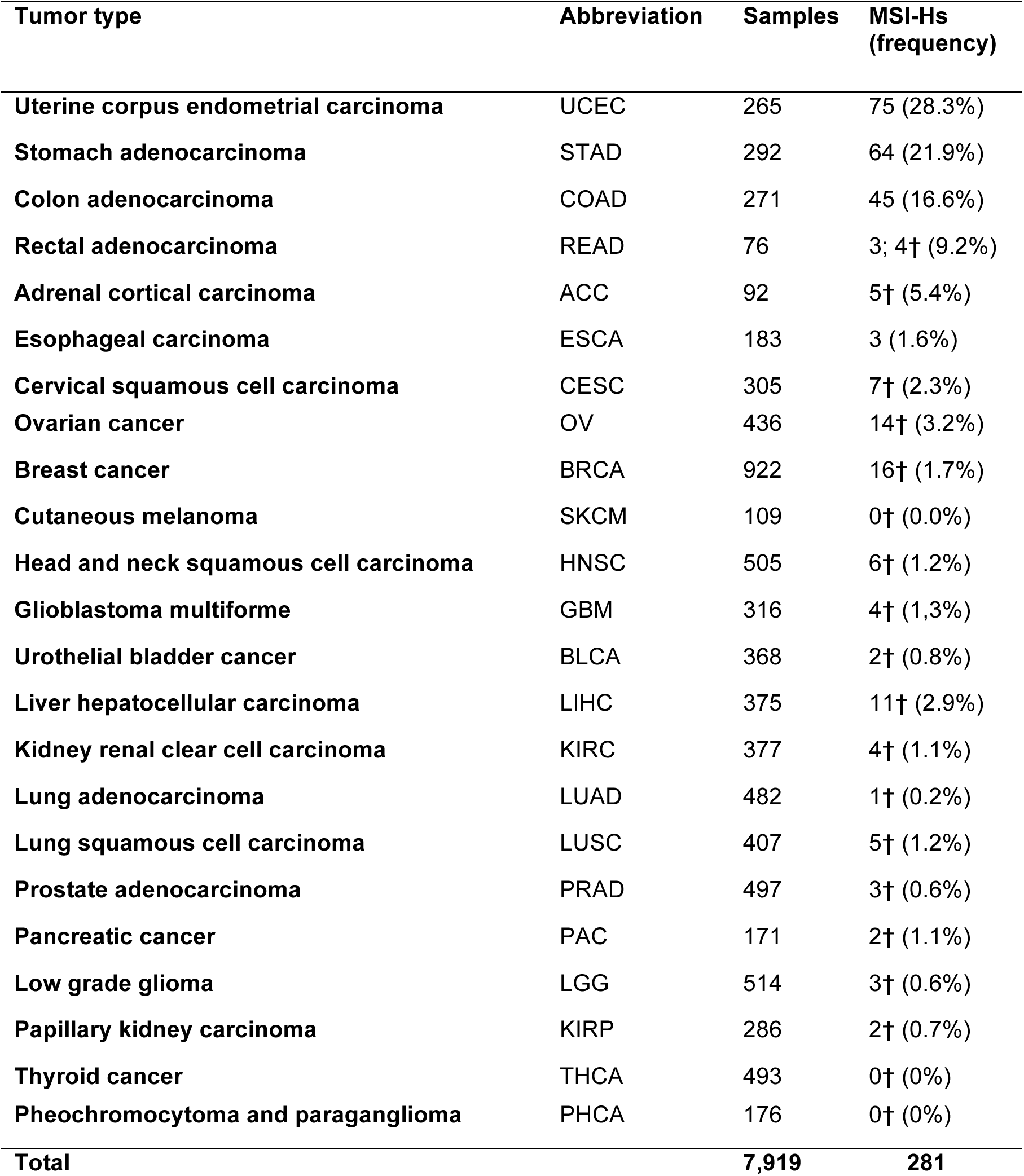
Exome tumors utilized to profile MSI. The Abbreviation column indicates the cancer type abbreviations used throughout the manuscript. The number of cases predicted as MSI-H at a confidence level of 0.75 is indicated with †.

| **Tumor type** | **Abbreviation** | **Samples** | **MSI-Hs**<br>**(frequency)** |
| --- | --- | --- | --- |
| **Uterine corpus endometrial carcinoma** | UCEC | 265 | 75 (28.3%) |
| **Stomach adenocarcinoma** | STAD | 292 | 64 (21.9%) |
| **Colon adenocarcinoma** | COAD | 271 | 45 (16.6%) |
| **Rectal adenocarcinoma** | READ | 76 | 3; 4† (9.2%) |
| **Adrenal cortical carcinoma** | ACC | 92 | 5† (5.4%) |
| **Esophageal carcinoma** | ESCA | 183 | 3 (1.6%) |
| **Cervical squamous cell carcinoma** | CESC | 305 | 7† (2.3%) |
| **Ovarian cancer** | OV | 436 | 14† (3.2%) |
| **Breast cancer** | BRCA | 922 | 16† (1.7%) |
| **Cutaneous melanoma** | SKCM | 109 | 0† (0.0%) |
| **Head and neck squamous cell carcinoma** | HNSC | 505 | 6† (1.2%) |
| **Glioblastoma multiforme** | GBM | 316 | 4† (1.3%) |
| **Urothelial bladder cancer** | BLCA | 368 | 2† (0.8%) |
| **Liver hepatocellular carcinoma** | LIHC | 375 | 11† (2.9%) |
| **Kidney renal clear cell carcinoma** | KIRC | 377 | 4† (1.1%) |
| **Lung adenocarcinoma** | LUAD | 482 | 1† (0.2%) |
| **Lung squamous cell carcinoma** | LUSC | 407 | 5† (1.2%) |
| **Prostate adenocarcinoma** | PRAD | 497 | 3† (0.6%) |
| **Pancreatic cancer** | PAC | 171 | 2† (1.1%) |
| **Low grade glioma** | LGG | 514 | 3† (0.6%) |
| **Papillary kidney carcinoma** | KIRP | 286 | 2† (0.7%) |
| **Thyroid cancer** | THCA | 493 | 0† (0%) |
| **Pheochromocytoma and paraganglioma** | PHCA | 176 | 0† (0%) |
| **Total** |  | **7,919** | **281** |

**Figure 1.**
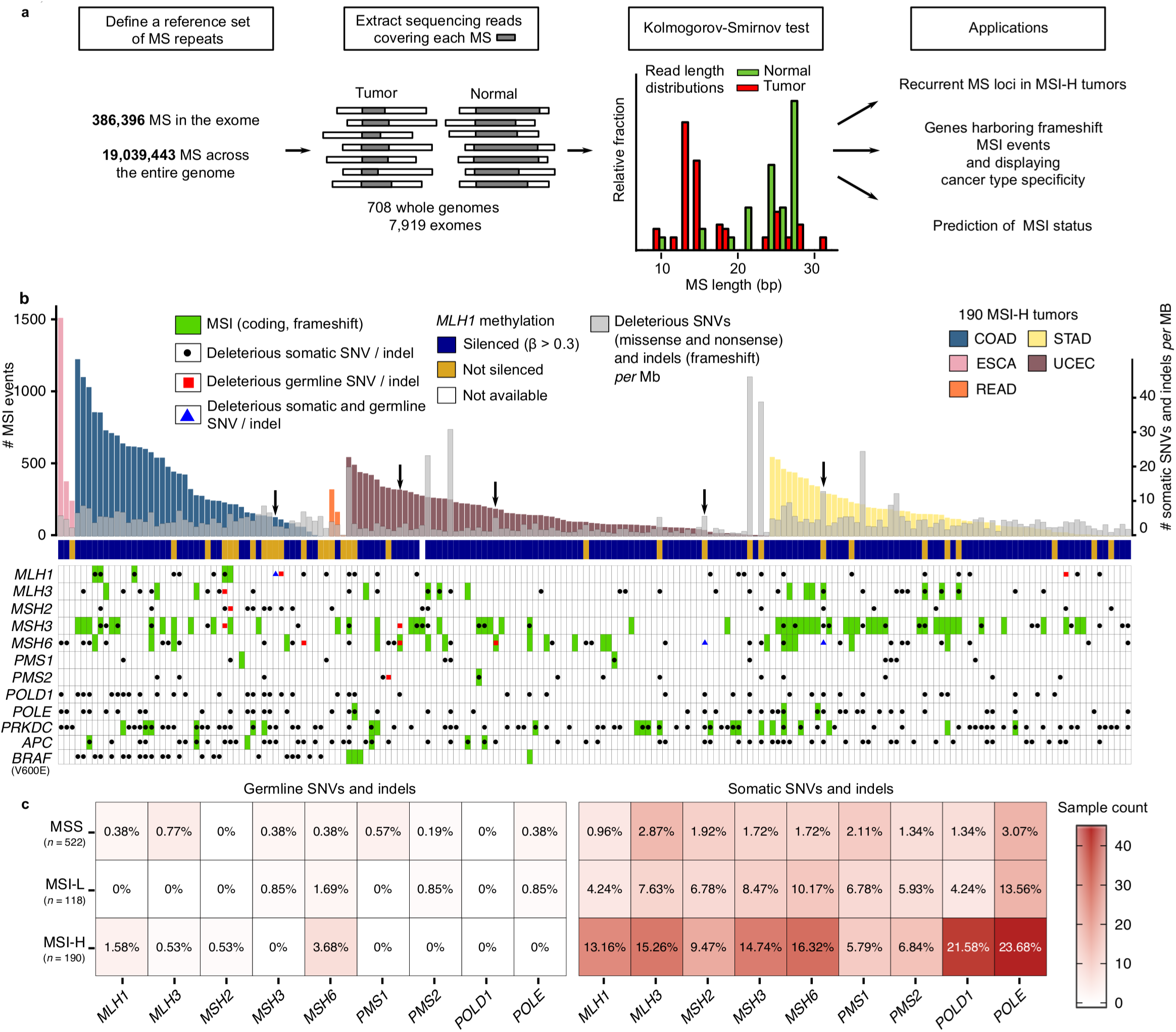
Schematic overview of the MSI calling pipeline. (**a**) A reference set of exonic and genome-wide MS repeats was assembled from the human reference genome hg19. The sequencing reads spanning each MS repeat and at least 2 base pairs at each flanking side were extracted from the tumor and normal BAM files. This process was repeated for all MS repeats in the reference sets across all pairs of matched normal-tumor samples. The Kolmogorov-Smirnov test was used to evaluate whether the read length distributions from the normal and tumor samples differed significantly (False Discovery Rate < 0.05). The exonic and genome-wide MSI calls served to identify MS loci recurrently altered by MSI in MSI-H tumors, discover frequent frameshift mutations and to predict MSI status. (**b**) Landscape of somatic MSI in MSI-H tumors. MSI events (frameshift and in-frame), deleterious SNV (missense, nonsense, and splice site) and indel (frameshift) rates in 190 MSI-H exomes. Samples harboring hypermethylation of the *MLH1* promoter are denoted by blue squares. Deleterious germline and somatic mutations (i.e., missense, nonsense, splice site and frameshift) are depicted in black and red, respectively, whereas frameshfit MSI events are shown in green. Black arrows mark patients with germline and somatic mutations in MMR genes. (**c**) Germline and somatic mutations in MMR genes, *POLE* and *POLD1* in MSS, MSI-L and MSI-H tumors. The heatmap and the cell labels report the number and percentage of samples in each category harboring mutations, respectively.

We first investigated five tumor types (COAD: colon adenocarcinomas, ESCA: esophageal carcinoma, READ: rectal adenocarcinoma, STAD: stomach adenocarcinoma and UCEC: uterine corpus endometrial carcinoma) for which the MSI status was determined by TCGA using capillary sequencing-based fragment length assay (Supplementary Table 1)^16–18^. These five tumor types have been recognized as MSI-prone and comprise the majority of MSI events amongst the samples we examined (44,462 MSI events in these five tumor types, *n*=904, versus 29,659 events in the remaining cases, *n*=7,015). Figure 1b shows the abundance of MSI events across the 190 MSI-H cases in these five tumor types (see Supplementary Figs. 1 and 2a for the remaining 118 MSI-L (MSI-Low) and 522 MSS tumor genomes in these 5 tumor types, and Supplementary Fig. 2b and Supplementary Table 1 for all tumor types). Our analysis highlights that MSI mutations represent a continuous rather than a dichotomous phenotype. The figure also shows a pronounced variability in the number of MSI events across MSI-H cases and across cancer types, indicating substantial intra-and inter-tumor type heterogeneity in the genomic impact of MSI (Supplementary Fig. 1 and Table 1). For example, we note that 7% and 17% of the MSI-H tumors have fewer than 10 and 50 detected MSI events, respectively, including one COAD MSI-H tumor without any exonic MSI events, while others have several hundred (exome sequencing data covers some neighboring non-exonic elements including small fractions of UTRs and introns; our term ‘exonic’ includes those spill-over regions).

**Figure 2.**
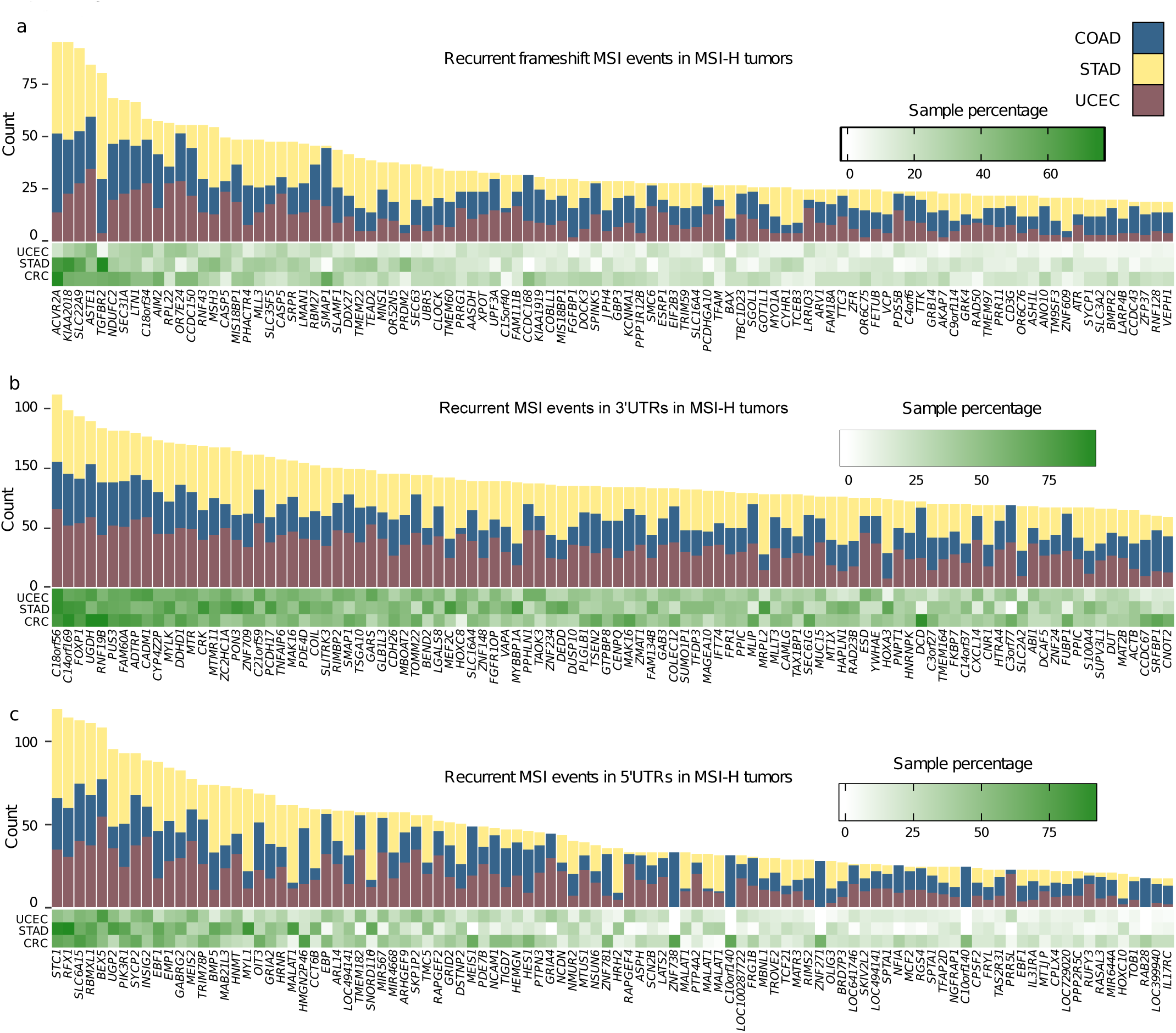
MS loci recurrently altered by MSI. (**a**) Coding MSI loci recurrently targeted by frameshift MSI in COAD, STAD and UCEC MSI-H tumors. The heatmap shows the fraction of COAD, STAD and UCEC MSI-H tumors containing frameshift MSI events in MS loci located within the coding sequence of the genes indicated on the *x*-axis. The total count of frameshift MSI events at these loci is depicted in the above barplot. The full list of MS loci recurrently altered by frameshift MSI is given in Supplementary Table S4. Similarly shown for genes with frequent 3' UTR (**b**) and 5’ UTR (**c**) MSI events in 3 MSI-prone tumor types.

To discriminate clonal MSI events from subclonal ones, we examined the CCF (cancer cell fraction) using the allele counts of sequencing reads containing the wildtype and mutant alleles^19^. The majority (87%) of the MSI events (32,327 events observed in 146 MSI-H cases for which tumor purity and absolute and local copy number data could be computed^20^) are clonal (CCF > 0.95, Supplementary Fig. 3a). We also observe that the relative abundance of MSI events is largely comparable across tumor stages (I-IV). This suggests that the mutational burden that tumors can tolerate may be similar across early and late tumor stages (Supplementary Fig. 3b). We note that subclonal MSI events may be under-represented relative to clonal events in late stages (Stage II - IV) compared to Stage I (Supplementary Fig. 3b), which would suggest that these genomes are more subject to clonal sweeps during the late stages.

**Figure 3.**
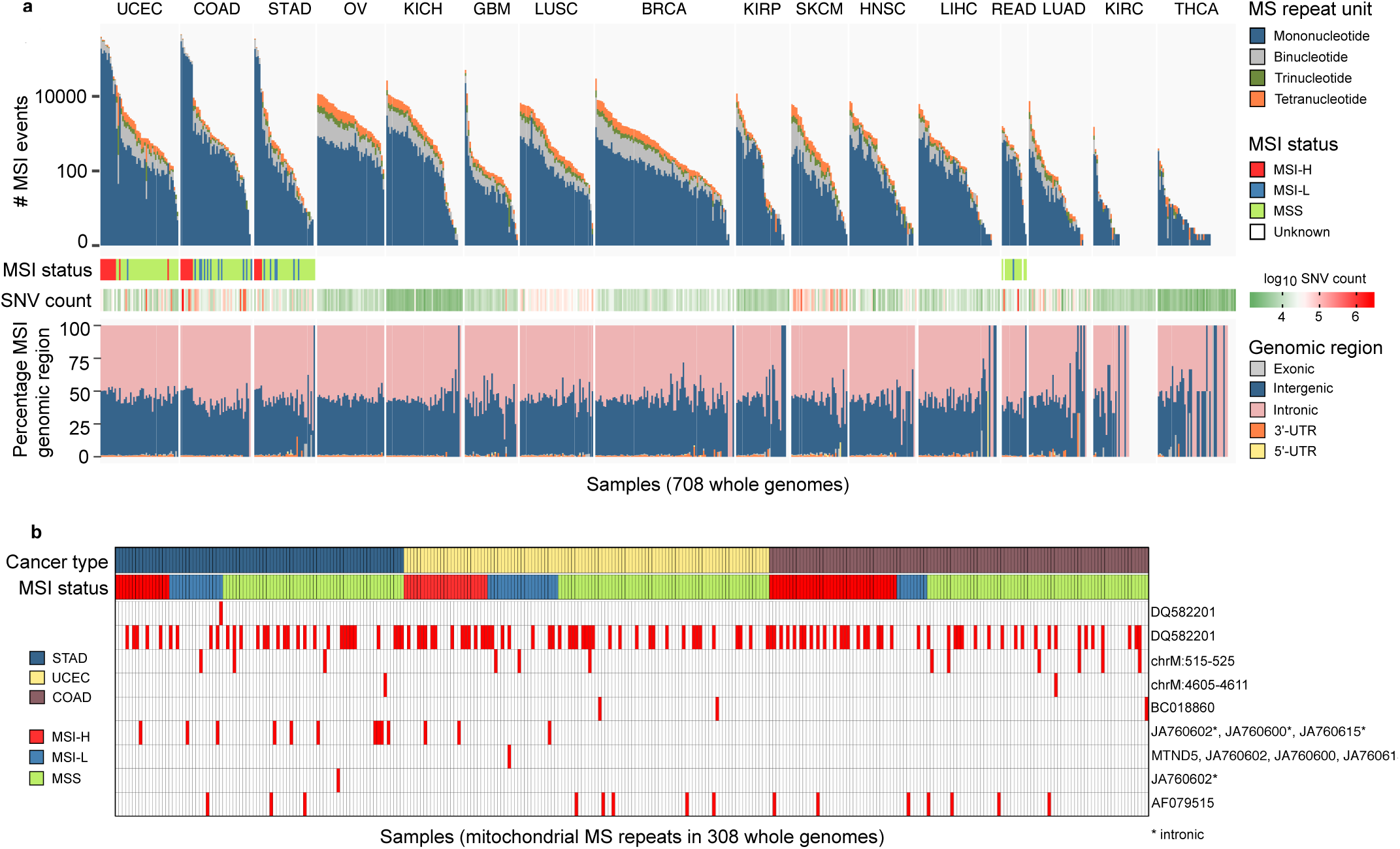
Pan-cancer landscape of genome-wide MSI. (**a**) The first panel shows the number of MSI events across 708 whole genomes, stratified by the length of the repeat unit. The second and third panels report the MSI status and the total count of SNVs, respectively. The fourth panel shows the distribution of MSI events across the genome. (**b**) Landscape of MSI in mitochondrial DNA across 308 COAD, STAD and UCEC low-pass whole genomes. MSI events, including frameshift and in-frame mutations, are shown in red.

Next, we identified genes with recurrent MSI events using MutSigCV^21^. The genes displaying significant enrichment for coding MSI (FDR < 0.05) along with their level of significance across three tumor types are shown in Suplementary Fig. 4. Pathway analysis reveals that transmembrane/TGFβ, response to cellular stress/DNA damage and chromosome/M-phase-related molecular functions are significantly enriched in genes harboring recurrent MSI in COAD, STAD and UCEC cases, respectively (Supplementary Table 2; P < 0.01).

**Figure 4.**
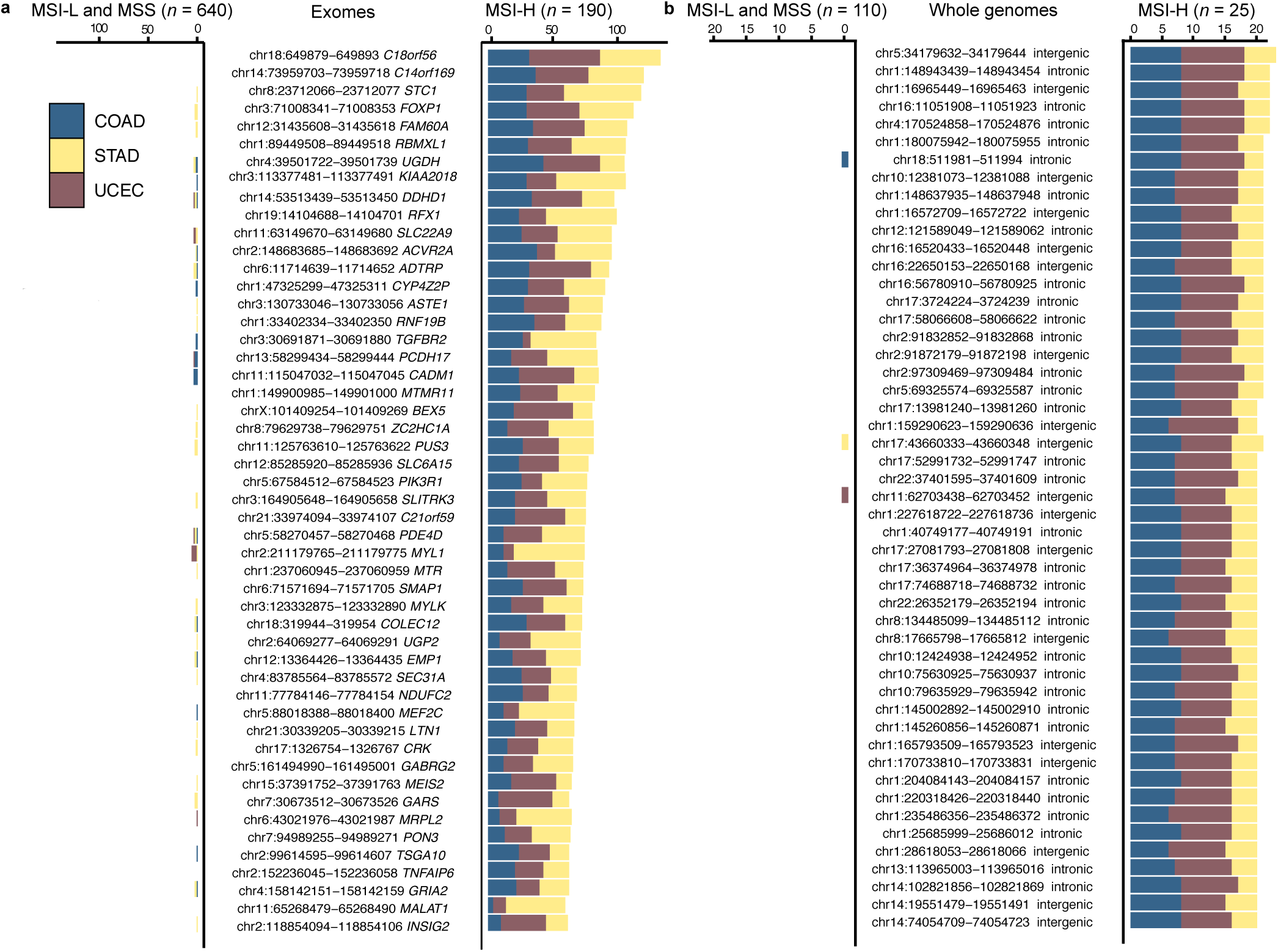
MS repeats recurrently altered by MSI in MSI-H tumors. (**a**) The barplots report the number of COAD, STAD and UCEC tumors harboring MSI events at the loci indicated in the central panel. This analysis examined 190 MSI-H, 118 MSI-L and 522 MSS exomes. (**b**) The recurrence analysis was extended to 25 MSI-H, 19 MSI-L and 105 MSS whole-genomes. Genomic coordinates in (**a**) and (**b**) indicate the location of the MSI repeats in the hg19 assembly of the human genome.

### The mutational landscape of DNA repair pathways

The rates of deleterious mutations (i.e., missense, nonsense and splicing site SNVs, as well as frameshift indels) and frameshift MSI events for *MLH1, MLH3, MSH2, MSH3, MSH6, PMS1, PMS2, POLE, POLD1, PRKDC, APC* and *BRAF* (p.V600E) are shown in Figure 1b. Among these genes, selected on the basis of their known implication in MSI, DNA repair and colorectal cancer, MSI frameshift events represent a major source for the inactivation of *MSH3* (85%) and *MSH6* (85%). In contrast, deleterious SNV mutations more frequently contribute to the loss of function of *POLD1* and *POLE* (27% and 23% of cases, respectively). We next examined the patterns of frameshift MSI events across MSI-prone tumors. We selected a set of 151 genes^22^ involved in several DNA repair pathways, including non-homologous end joining (NHEJ), homologous recombination (HR), base excision (BER), RecQ helicase-Like (RECQ), translesion synthesis (TLS), and ataxia telangiectasia mutated (ATM)^22^. We find that COAD samples harboring a large number of MSI events (>500 in our samples) are enriched for *MLH1* promoter hypermethylation (Figure 1b), as previously reported for this tumor type^15^. The genes most frequently targeted by MSI are *RAD50* (16% of MSI-H tumors), *ATR* (15%) and *RBBP8* (10%) (Supplementary Table 3 and Supplementary Fig. 5a).

**Figure 5.**
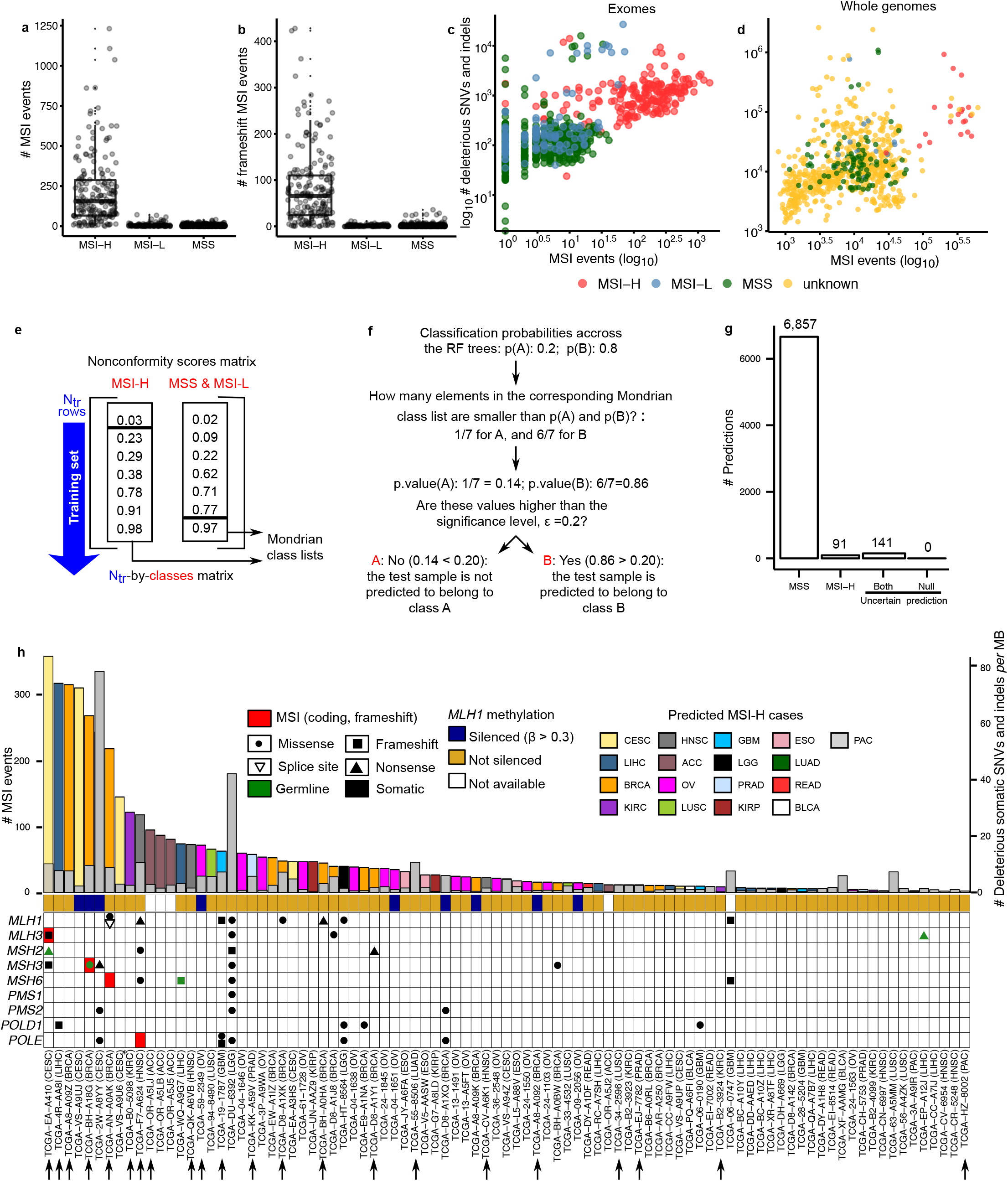
Distribution of the number of MSI and prediction of MSI status. Distribution of the number of MSI (**a**) and frameshift MSI events (**b**) in MSI-H and MSS (also including MSI-L) tumors. Correlation between the number of SNV and MSI events in exomes (**c**), and whole genomes (**d**). Prediction of MSI status from exome-sequencing data using conformal prediction and Random Forest models (**e**). Initially, we used 5-fold cross-validation to calculate predictions for all training examples. The fraction of trees in the forest voting for each class was recorded, and subsequently sorted in increasing order to define one Mondrian class list *per* category. (**f**) The model which was trained on all training data was applied to 7,089 exomes. For each of these samples, the algorithm recorded the fraction of trees voting for each class. The *P* value for each class was calculated as the number of elements in the corresponding Mondrian class list higher than the vote for that class (e.g. 6 out of 7 in the toy example depicted in Fig. 5a) divided by the number of elements in that list. If the *P* value for a given class is above the tolerated error, ε, the sample is predicted to belong to that category. The confidence level (1-ε) indicates the minimum fraction of predictions that are correct. (**g**) Number of samples predicted as MSI-H, MSS and uncertain (both: in cases when the classifier does not have enough power to confidently assign a single category; none: in cases when the samples that are outside the applicability domain of the model). Here, the confidence level was set to 0.75. (**h**) Landscape of MSI for the 91 exomes predicted as MSI-H at a confidence level of 0.75. Samples predicted to be MSI-H at a confidence level of 0.80 are marked with black arrows.

We have also examined the impact of germline mutations in the MSI-H cases. We observe that 4 COAD (9%), 4 UCEC (5%) and 2 STAD (3%) patients harbor deleterious germline mutations in MMR genes. Of these, at least 5 patients may have acquired the MSI-H phenotypes due to biallelic inactivation of MMR genes, where the inherited germline mutations of MMR genes are complemented with somatically acquired mutations of the corresponding genes. One COAD sample harbored germline and somatic mutations in *MLH1*; 1 STAD and 3 UCEC cases harbor germline and somatic mutations in *MSH6.* Overall, germline mutations in MMR genes, *POLE* and *POLD1* are consistently more prevalent in MSI-H patients compared to MSS cases (Fig. 1c and Supplementary Fig. 6). This frequency of germline mutation carriers in MMR genes may be underestimated since we have applied stringent filtering criteria for our germline calls (Online Methods) to account for the uncertain pathogenicity of missense mutations^23^, as well as the technical challenges in identifying mutations in *PMS2* with multiple copies of its pseudogenes in the genome^24^.

Although it is difficult to pinpoint the genomic events initiating MMR deficiency, it is likely that truncating mutations in various MMR genes in addition to the hypermethylation of *MLH1* may shape MSI-H genomes, leading to further accumulation of mutations in genes of the DNA repair pathway. To investigate the downstream impact of somatic alterations in MMR genes and proofreading DNA polymerases, we examined the correlation between gene expression and promoter methylation, DNA copy numbers, somatic SNVs and indels and MSI events (Supplementary Fig. 7). For *MLH1*, only the DNA methylation level is associated with gene expression levels (r = -0.79; Pearson correlation), consistent with a previous report^3^. No apparent relationship between promoter methylation and gene expression is observed for the other genes examined. Other than *MLH1*, the most common genomic events that show association with gene expression (*P* < 0.05; Mann-Whitney test) are the truncating SNVs and frameshift MSI events (*MLH3*, *MSH2*, *MSH3*, *MSH6*, *PMS1* and *POLD1*), suggesting that these somatic events are responsible for the under-expression of the genes in which they are located and may also play a major role in the development of the MSI-H phenotype. The association between DNA copy number and gene expression (r > 0.2; Pearson correlation) is observed for *MSH2* and *POLD1*. We do not observe any significant association between gene expression and germline truncating mutations.

### Cancer-type specificity in loci frequently targeted in frameshift MSI events

We investigated the frequency of frameshift MSI events in 130 cancer-related genes^25^ across the MSI-prone tumors. Tumor-type specificity of frameshift MSI is evident for some well-known targets of MSI, such as *ACVR2A* (52% of MSI-H tumors) and *TGFBR2* (44%) (enriched in both COAD and STAD; *P* < 0.05, one-tailed Fisher’s exact test) as well as *RPL22* (31%), *RNF43* (31%), *MLL3* (27%), *PRDM2* (21%), *JAK1* (16%) and *APC* (3%) (Supplementary Fig. 5b and Supplementary Table 4)^26–28^. For instance, frameshift MSI events are present in *TGFBR2* for 26/45 (58%) of COAD and 51/64 (80%) of STAD but only in 4/75 (5%) of UCEC cases, suggesting that certain tumor types or tumor environments are favorable for the occurrence of particular MSI events. Given the nature of frameshift MSI events in coding regions (i.e., protein-truncating and thus inactivating), the absence of MSI in well-known oncogenes such as *BRAF* likely represents the pressures of negative selection in the context of the MSI-H phenotype. Instead, *BRAF* V600E mutations are observed in 22 of the 45 (49%) MSI-H COAD tumors, but only 4 (2%) frameshift MSI events are observed within the gene.

To uncover other MS loci frequently targeted by frameshift MSI mutations, we first ranked MS loci according to the recurrence level of frameshift MSI events in COAD, STAD and UCEC MSI-H tumors. This analysis resulted in 16,812 frameshift MSI events across a set of 6,441 coding MS loci spanning 4,898 genes, several of which display cancer type specificity (Fig. 2a and Supplementary Table 5). The most recurrent frameshift MSI events are found *in ACVR2A* (51.6% of the tumors)*, KIAA2018 (*51%), *SLC22A9 (*50%), *ASTE1* (45%*), TGFBR2* (44%), *NDUFC2*(36%), *LTN1* (36%) and *SEC31A* (36%). Frameshift MSI events often display significant tumor type specificity, e.g., *MLL3, PRDM2, C9orf114, BAX and OR7E24* are enriched in STAD*, JAK1, TFAM* and *SMC6* are enriched in UCEC, whereas *SEC31A, C18orf34, NDUFC2, KIAA1919* and *CCDC168* among others, are enriched in COAD (*P* < 0.05, one-tailed Fisher’s exact test). Among low-frequency MSI events, *SMAP1*, *CCDC168* and *SPINK5* harbor frameshift mutations in COAD and UCEC but not in STAD tumors.

By analyzing the frequency of MSI events in untranslated regions, we found that MS loci within the 3’ UTR region of *C18orf56*, *C14orf169*, *FOXP1*, *UGDH, RNF19B, PUS3* and *FAM60A* as well as the 5’ UTR region of *STC1, RBMXL1, RFX1, BEX5* and *SLC6A15* are recurrently altered by MSI across MSI-H cases (Fig. 2b,c and Supplementary Table 6,7). Other MS loci display marked cancer-type specificity, e.g., MSI events within the 5’ UTR region of *ZNF738*, *C10orf140, ZNF271 and RAB28* are specific for COAD tumors, whereas *EBP, TMEM182, MIR567* and *MEIS1* are depleted for MSI in STAD tumors. Supplementary Table 8 reports the enrichment of frameshift, 3' and 5' UTR MSI events in COAD, STAD and UCEC.

To obtain a comprehensive MSI landscape on a pan-cancer scale, we next extended our analysis to all exomes, irrespective of their status as MSI-H, MSI-L or MSS. We observe frameshift MSI events in 8,011 MS loci, of which 51 are altered in more than 50 samples (Supplementary Fig. 8 and Supplementary Table 9). *ACVR2A, TGFBR2, KIAA2018, ASTE1* and *SLC22A9* frequently harbor MSI events in STAD and COAD, whereas several other genes are mostly specific to a single tumor type. For instance, *FAM129A, GMIP* and *NEK3* are altered in 107 (12%), 93 (10%) and 53 (6%) BRCA (breast cancer) tumors, respectively; *ALPK2* and *DPYSL2* are altered by frameshift MSI in 73 (17%) and 59 (14%) OV (ovarian serous cystadenocarcinoma) patients, respectively, but only 62 times in the remaining samples; and *ABT1* and *SLC22A24* are altered in 19.6% and 14% of ACC (adrenocortical carcinoma) tumors. Although the cancer-related roles of these novel MSI targets are largely unknown, it has been shown that siRNA-mediated inhibition of *ALPK2* inhibits apoptosis,^29^ suggesting that the functional implication of these novel, recurrent MSI events warrants further investigation.

### A genome-wide mutational spectrum of MSI

Despite the plethora of studies on genome-wide somatic variation, the landscape of non-coding MSI in cancer remains uncharted territory. Most of the non-coding MSI events are neutral passengers, but those occurring in regulatory elements (see next section) can function as cancer drivers, similar to somatic SNVs in enhancer regions that have been shown to play a role in tumorigenesis.^30,31^ To profile the distribution of MSI events genome-wide, we leveraged 708 whole-genome (mean coverage: 55X) pairs of tumor and matched non-neoplastic samples spanning 16 cancer types (Fig. 3 and Supplementary Table 10). The number of MSI events in MSI-H tumors differs significantly from that in MSI-L (*P* = 6.25 x 10^-11^, Kolmogorov-Smirnov) and MSS (*P* = 4.01 x 10^-15^, Kolmogorov-Smirnov) tumors, whereas the numbers are comparable between MSI-L and MSS cases (*P* = 0.17, Kolmogorov-Smirnov). As shown in Fig. 3a, when samples are ordered by decreasing number of MSI events within each tumor type, the decrease is gradual, confirming that MSI represents a continuous rather than a dichotomous phenotype.

The numbers of exonic MSI calls between the exome and whole-genome datasets show a high correlation (r = 0.90, *P* < 10^-15^, Pearson correlation for 531 cases profiled on both platforms; Supplementary Fig. 9). However, many MSI events are missed in whole-genome data due to their lower coverage, with only 32% of the calls from the exomes captured on the genomes (based on 23 pairs with at least 50 MSI events in exome data). On the other hand, 13% of the exonic MSI events identified in whole-genome data are missed in the exome-based calls, since the target capture regions do not include many exonic MS repeats.

To further analyze the influence of read depth on MSI detection sensitivity, we performed subsampling analysis, using a sample with a high number of MSI events and coverage (TCGA-AD-A5EJ; tumor at 82X, matched normal at 42X) (Supplementary Figs. 10-11). We find that the number of MSI events recovered decreases substantially when the coverage is reduced to 20-30X. However, we do not see any clear relationship between the number of MSI events and coverage above that range (Supplementary Fig. 11), suggesting that sequencing coverage is not a major factor, especially given that most MSI events are clonal as we described earlier. We also examine the correlation between the number of MSI events and tumor purity (Supplementary Fig. 12).^20^ The number of MSI events identified in high purity samples spans five orders of magnitude, whereas we systematically detect fewer than a thousand MSI events in the case of low-purity samples. However, the number of low purity samples (e.g., <0.6) in our set is small, meaning that the impact of coverage due to variation in purity does not have a substantial impact on our analysis.

The genome-wide density of MSI events along the chromosomes does not show statistically significant correlation with SNV density, regardless of the bin size (100kb - 10Mb bins; Supplementary Fig. 13). We have previously used exome sequencing to identify a portion of MSI events in UTRs^15^. With whole-genome data, the number of MSI calls in 3’ UTRs is substantially larger (Supplementary Fig. S9b-d). This allowed us to find that MSI events are enriched in 3’ UTRs for MSI events in 21 out of 25 MSI-H cases (84%), whereas they are depleted in 5’ UTRs and coding regions in 24 out of 25 MSI-H cases. In MSS tumors, only 3 out of 105 (3%) show enrichment of MSI in 3’ UTR regions, whereas 42 (46%) show depletion of MSI events in coding regions (*P* < 0.05; one-tailed Fisher’s exact test). Overall, these results suggest that MSI events in 3’ UTRs may be under positive selection in MSI-H tumors. The shortening of 3' UTRs in cancer cells is known to increase the stability of transcripts and thus the translational level of oncogenes^32^. The frequent MSI events in 3' UTRs may have similar functional consequences (e.g., the loss of miRNA-mediated regulation), although they often result in down-regulation of their corresponding genes. ^33^

We find that 62% of the tumors harbored more than 100 MSI events genome-wide, including samples from all 16 cancer types examined. We again observe a substantial level of intra-tumor type heterogeneity in MSI abundance, with the number of MSI events varying up to 5 orders of magnitude. The presence of noncoding MSI events genome-wide in tumor types beyond the MSI-prone cases is notable. For instance, we observe that 81% and 53% of OV and KICH (kidney chromophobe) samples, harbor >1,000 MSI events, exceeding the previously reported MSI-H frequencies of OV (12%) and overall kidney cancers (6%)^34,35^. Elevated microsatellite alterations at selected tetranucleotides (EMAST) has been observed in non-MSI-prone tumor types such as lung, head and neck cancers as well as melanoma^36^, but the current markers for EMAST could not be captured with the short reads in our data due to their size.

To investigate the relationship between epigenetic features and the genome-wide distribution of MSI events, we selected the 50 genomes displaying the highest MSI rates and compared their MSI density with the 25-state chromatin state map based on 12 epigenetic marks across 127 epigenomes^37^ (Online Methods; Supplementary Figs. 14-17). For best estimates, the chromatin state map from the most ‘similar’ tissue type was used for each tumor type. Our analysis reveals significant enrichment for MSI events in actively-transcribed regions, promoters and enhancers in most MSI-H genomes (two-tailed Fisher’s exact test, *P* < 0.05; Supplementary Fig. 18). On the other hand, inactive regions, including constitutive heterochromatin, repressed Polycomb, bivalent promoters and quiescent chromatin, are significantly depleted for MSI overall. Taken together, these results show the over-representation of MSI in functionally important, typically open-chromatin regions, extending our previous results based on seven colorectal and UCEC tumors^15^.

### MSI events in the mitochondrial genome

To obtain a mitochondrial MSI landscape, we analyzed TCGA low-coverage (6-8X) whole-genome sequencing data that we have generated (the sample size in this dataset was larger than that of the high coverage dataset). Due to their high copy number, the mitochondrial DNA had a median coverage of >800X even in the low-coverage data^38^. We applied our MSI discovery pipeline to a set of 31 mitochondrial MS loci (Supplementary Table 11) across 308 cancer genomes (COAD, STAD and UCEC) (Fig. 3b and Supplementary Table 12). The most recurrent MSI event is observed in *DQ582201*, a polyC mononucleotide repeat (115 MSI events, 37% of tumors); the second most recurrent event is on the exon regions of *AF079515* (15 MSI events, 5% of tumors). The instability of *DQ582201* has been reported in several cancer types.^39^ The majority of mitochondrial MS loci (22 out of 31 MS loci; 71%), however, do not contain MSI events in any of the tumors examined, suggesting that mitochondrial MSI is not widespread compared to nuclear MSI. Moreover, mitochondrial MSI events are not associated with the MSI status of the tumor: 43 (54%), 22 (48%) and 98 (54%) mitochondrial MSI events are observed in MSI-H, MSI-L and MSS genomes examined, respectively. The relationship between the nuclear MSI and mitochondrial MSI has been controversial^39^, but our mitochondrial genome-wide MSI examination suggests that these two events are not correlated.

### Defining a panel of hypermutable MS loci enriched in MSI-H cases

We focused on frameshift MSI events earlier to understand the functional impact of coding MSI in MSI-prone tumors. Here, we sought to uncover mutational patterns discriminative of MSI-H status across exonic and noncoding MS loci, as well as to identify hyper-mutable MS loci across the entire genome. We first ranked recurrently-targeted MS loci by their specificity for MSI-H tumors using exome sequencing data from COAD, STAD and UCEC tumors. Our analysis yielded a catalogue of MS loci specific to MSI-H tumors (Fig. 4a and Supplementary Table 13). We find that several of these MS loci lie within genes prone to frameshift MSI events, such as *KIAA2018*, *ACVR2A* or *ASTE1*^15^. In contrast to frameshift and 3’/5’ UTR MSI events (Fig. 2), few MSI events enriched in MSI-H and depleted in MSS/MSI-L cases display cancer-type specificity, implying that there are commonalities in the molecular mechanisms underlying MSI at these loci across the three cancer types.

Given that most MS loci lie within the non-coding genome, we also extended our recurrence analysis to whole-genomes by utilizing sequencing data from 25 MSI-H, 19 MSI-L and 105 MSS tumors. We discovered a set of intronic and intergenic MS repeats recurrently targeted by MSI in MSI-H cases (Fig. 4b and Supplementary Table 14). Perhaps not surprisingly given the larger list of candidate MS, these non-coding loci are more specific to MSI-H than the best exonic loci are (cf. Fig. 4a), containing MSI events in nearly all of the MSI-H tumors and almost none in MSI-L or MSS samples. These inquiries have yielded a collection of coding and non-coding MS loci recurrently targeted by MSI in MSI-H tumors, which provide a foundation to refine and extend the set of markers employed for MSI-H categorization.

### Prediction of MSI status from exome-sequencing data

The total numbers of MSI and frameshift MSI events are significantly higher in MSI-H tumors than in MSI-L or MSS tumors (*P* < 10^-15^; Kolmogorov-Smirnov test; Figs. 5a,b). The number of SNV and MSI events exhibit moderate to low correlation in MSI-H (r = 0.32, *P* = 6.15 x 10^-6^ in exomes, and r = 0.35, *P* =0.09 in whole genomes; Pearson correlation; Figs. 5c,d), MSI-L (r = 0.10, *P* = 0.68, Pearson correlation) and MSS (r = -0.06, *P* = 0.56, Pearson correlation) tumors. To test whether our MSI calls can be used to distinguish between MSI-H and MSS cases, we built Random Forest^40^ classification models. Each tumor was encoded by a vector recording the presence or absence of MSI events across MS loci, as well as the total number of MSI events (Online Methods). Models built on a limited set of learning examples (i.e., only those 190 MS-H, 118 MSI-L and 533 MSS tumors with MSI status annotations) are likely to possess limited predictive power on external data. Thus, we included conformal prediction^41^ in our modeling pipeline to provide confidence estimates for individual predictions (Fig. 5e,f; Online Methods). Briefly, conformal prediction evaluates the similarity (i.e., conformance) between the new samples and the training data. The output represents the probability that the new sample is either MSI-H, MSS, or uncertain (in the case of the new samples being outside the applicability domain of the model), given a user-defined tolerance level, which sets the maximum allowable fraction of erroneous predictions. Our 10-fold cross-validation (CV) showed high accuracy of the models produced (sensitivity: 92%; specificity: 99%). Comparable results were obtained in Leave-One-Out CV (sensitivity: 93%; specificity: 99%), indicating that the MSI events detected using whole-exome data convey enough predictive signal for MSI categorization.

By applying the prediction model to 7,089 exomes from 17 cancer types not commonly tested for MSI status, we identified 91 additional MSI-H cases using a confidence level of 0.75, 22 of which were identified at confidence level of 0.80 (Figs. 5g,h and Supplementary Table 15). Among the 91 cases, the most frequent are BRCA (16), OV (14) and LIHC (liver hepatocellular carcinoma; 11). Our estimated MSI-H rate for ovarian cancer is 3.2%, significantly lower than that reported previously (10%)^42^; for HNSC (head and neck squamous cell carcinoma) and CESC (cervical cancer), our estimated MSI-H rates are 1.2% and 2.3% whereas the reported rates in the literature are 3% and 7%.^8^ The frequencies generated for the other non-MSI-prone cancer types were mostly in agreement with the reported numbers in the literature.^8^ For example, our estimated MSI-H frequencies for PRAD (prostate adenocarcinoma), LUAD (lung adenocarcinoma) and LUSC (lung squamous cell carcinoma) are 0.6%, 0.2% and 1.2%, respectively, which are comparable to the frequencies of 1% and 0-2% reported for prostate and for lung cancers, respectively^8^. We note that the differences in the rates may be due to the small sample sizes used in the literature for some tumor types^8^, differences in the characteristics of the cohorts (e.g., tumor stage) and tumor-type specific features that were missed in our model. We did not identify any MSI-H cases among THCA (papillary thyroid carcinoma; *n*=493), PHCA (pheochromocytoma; *n*=179) and SKCM (skin cutaneous melanoma; *n*=109) tumors. Overall, the frequency of MSI-H cases in non-MSI-prone cancer types was found to be 1.3%, significantly lower than the 14% we observed in UCEC, STAD, COAD, READ and ESCA tumors. Consistent with our analyses of COAD, READ, STAD, ESCA and UCEC MSI-H tumors (Fig. 1b), we found that the number of MSI events varied markedly across these newly identified MSI-H tumors (Fig. 5h). We detected 1,365 frameshift MSI events in the tumors predicted as MSI-H, with the most frequent incidences in *DPYSL2* (12 cases), *OR11G2* (9), *SLC22A9* (9) and *KIAA2018* (8), suggesting that the MSI events that recur in MSI-H cases (*cf.* Figure 2) constitute a mutational signature that is leveraged by the predictive model for MSI categorization. We find that 31 patients display somatic mutations in MMR genes, and 1 CESC (TCGA−EA−A410) and 2 LIHC (TCGA-WQ-A9G7 and TCGA-EP-A12J) cases harbor germline mutations in *MSH2*, *MSH6* and *MLH3*, respectively. Additionally, we observe that 1 BRCA patient (TCGA-BH-A18G) harbors a missense germline mutation predicted to be pathogenic with high confidence (Online Methods) and a somatic frameshift event in *MSH3*.

We also performed mutation signature analysis based on the mutation frequencies of 96 trinucleotide contexts^43^ (Supplementary Fig. 19). For the 91 MSI-H predicted cases, we confirm the mutation signatures characteristic of MSI-H cases, e.g., C>T transitions in (A/C/G)pCpN sequence contexts and C>A transversions at an CpCpN context, suggesting that the mutation signatures of predicted MSI-H cases are largely concordant with those of known MSI-H cases.

## Discussion

Our pan-cancer profiling of exome and whole-genome sequencing data for MSI has demonstrated that the number of MSI events varies profoundly within and across tumor types. Moreover, this variability is also observed within the MSI categories established by the Bethesda guidelines (MSI-H, MSI-L and MSS). These data therefore support the argument that microsatellite instability represents a continuous phenotype.

A joint analysis of MSI-H tumors from multiple cancer types has revealed that several DNA repair pathways other than MMR, including ATR, BER, HR and NHEJ, are altered by single-nucleotide and MS mutations. Moreover, we have uncovered new genes affected by frameshift MSI events in MSI-prone tumors as well as in tumor types not frequently affected by MSI (e.g., *FAM129A, GMIP* and *NEK3* in BRCA, and *DPYSL2 and ALPK2* in OV). Some of these genes have shown strong predictive power for MSI-H status (e.g., *ACVR2A* and *KIAA2018* for COAD, STAD and UCEC), whereas others display low recurrence and specificity for single cancer types (e.g., *SMAP1* for STAD) Along with the diverse molecular functions enriched for MSI events in these tumor types, our data reaffirm that some genes are particularly susceptible to MSI in certain cancer types^44^. Although some of their potential cancer-related roles have been identified^45^, the functional relationship between MSI and tumorigenesis as well as the similarity of molecular mechanisms establishing MSI phenotype across cancer types remain to be validated.

By classifying 7,089 patients into MSI-H and MSS categories using our MSI-based predictive model, we identified 91 new MSI-H cases from 16 different tumor types. According to our classification model, the frequency of MSI-H cases in MSI-prone tumors is roughly ten times larger than in other tumor types (14.5% vs 1.3%). In contrast to previous models based on SNVs or mononucleotide repeats,^46,47^ our modeling approach estimates the likelihood of prediction error for individual patients using a confidence level, which can be easily interpreted (e.g., a confidence level of 0.9 means that no less than 90% of the predictions should be correct).

Although the search space considered in our whole-genome and exome MSI reference sets is large (~19 million and 386,396 MS loci, respectively), it comprises only MS repeats of size 6-60 bp and up to tetrameric repeats. Although we are missing longer repeats as well as MS loci longer than 60 bp, our MSI calling pipeline captures the vast majority of MS loci (e.g., >99% of repeats in the final set of exome and genome reference MS are smaller than 40 bp).^15^ Nonetheless, MSI events in certain non-coding MS loci might have missed the threshold for significance due to low coverage, and we anticipate that the rates of MSI events presented here are likely to be under-estimates of the true rates. The sequencing data used in this study were obtained from a sequencing platform (Illumina) that is known to be robust in estimating the length of homopolymer runs,^48^ but a platform with longer reads will help in a more comprehensive identification of MSI target loci. A further increase in sample size, especially those with annotated MSI status to include in a better training set, will also increase the power to detect all relevant loci.

Of clinical relevance, we provide the largest catalogue available to date of coding and non-coding MS loci frequently altered across cancers. This repertoire has implications for MSI categorization and provides the groundwork for further investigation into the molecular mechanisms underlying MSI recurrence. For many cancer patients, the use of targeted panels containing only a subset of cancer-related genes is informative for personalized therapeutics. Such panels offer the advantage of very high sequencing depth (e.g., >1000X) to detect somatic mutations with low variant allele frequencies that would be missed by standard exome sequencing. The loci identified in our study, especially those in the non-coding regions, can be included in panel-based assays to serve as highly sensitive markers for MSI across multiple tumor types. This will avoid a separate test for MSI in MSI-prone tumors, and it will identify a small set of MSI patients in non-MSI-prone tumors in which MSI is rarely considered by clinicians. The potential benefit of such a panel is enormous, given the demonstrated efficacy of immunotherapy for the MSI cohort.^13^

## Online Methods

### Data sets

Exome and whole-genome tumor-normal pairs from the TCGA project were downloaded from CGHub (http://cghub.ucsc.edu). The reads were mostly 100 bp paired-end reads and were aligned to the NCBI build 37 (hg19) using BWA. The full list of samples is given in Supplementary Tables 1, 10 and 12. The MSI status (MSI-H, MSI-L and MSS) were downloaded from the GDAC (https://gdac.broadinstitute.org) website, whereas the methylation state of the *MLH1* promoter, gene expression and DNA copy number variation data were downloaded from the Genomics Data Commons Data Portal (https://gdc-portal.nci.nih.gov/) websites. MSI status was evaluated by the TCGA consortium for COAD, READ, ESCA, STAD and UCEC tumors using a panel of 4 mononucleotide repeats (BAT25, BAT26, BAT40 and TGFBRII) and 3 dinucleotide repeats (D2S123, D5S346 and D17S250), except for a subset of COAD/READ genomes evaluated by 5 mononucleotide markers (BAT25, BAT26, NR21, NR24 and MONO27)^49^. Tumors were classified as MSI-H (>40% of markers altered), MSI-L (<40% of markers altered) and MSS (no marker altered).

### Defining a reference set of MS repeats

To generate an exome-wide reference set of MS loci, we utilized the Sputnik algorithm^15^ to identify MS loci in the mRNA sequences of 39,496 RefSeq genes (USCS Genome Browser, hg19). We limited our analysis to mono-, di-, tri-and tetranucleotide MS loci of size 6-60 bp, which can be detected reliably with enough flanking sequences from 100 bp reads. We derived the reference set of MS repeats from RefSeq sequences as the target regions used by the TCGA are different across cancer types. MS repeats falling within splice sites were removed, as they have undetermined genomic coordinates or are redundant to multiple isoforms. The final reference set of exonic MS sites (Supplementary Table 16) comprised 386,396 loci (112,896 mono-, 63,162 di-, 132,117 tri-and 78,221 tetranucleotides). These included 154,590 coding, 50,598 5’-UTR and 181,208 3’-UTR MS loci, as annotated in the UCSC Genome Browser.

To produce a genome-wide reference set of MS loci (Supplementary Table 17), a total of 19,039,443 MS repeats were obtained using the Sputnik algorithm (chromosome 1 through Y) and categorized into five groups (coding, 248,100; 5’-UTR, 39,582; 3’-UTR, 166,111; intronic, 8,265,436; intergenic, 10,320,214). This MS set encompasses 7,404,614 mono-, 3,686,129, di-, 3,750,887 tri-and 4,197,813 tetranucleotides. We also utilized the Sputnik algorithm to build a reference set of mitochondrial MS loci from the hg19 mitochondrial DNA (mtDNA), which contained a total of 31 MS loci (10 mono-, 2 di-, 11 tri-and 8 tetranucleotides) (Supplementary Table 11).

### Detection of DNA slippage events

After filtering reads with low mapping quality, intra-read MS repeats were identified with the same method used to identify reference MS repeats and were intersected with the reference MS repeats. We note that the minimum size of intraread repeats detected was 5bp. Thus, reads spanning MS repeats contracted below 5bp were not considered. We required the 2 bp flanking sequences (both 5’ and 3’) of the intra-read MS repeats to be identical to those of matching reference repeats, thereby discounting truncated MS repeats. In each genome, the distribution of the allelic repeat length at each MS locus was obtained by collecting the lengths of all intra-read MS repeats mapped to that locus. We compared the distributions of MS lengths from tumor and matched normal genomes at each locus using the Kolmogorov-Smirnov statistic. An FDR (false discovery rate) of < 0.05 was used as a threshold for statistical significance, with a minimum of 5 tumor and 5 matched normal reads. We note that the number of MSI ‘‘events’’ refers to the absolute number of MSI counts per sample, whereas sample percentage refers to the percentage of samples from a given cancer type harboring MSI events at a particular MS locus. We distinguished MSI events at coding sequences into in-frame and frameshift events depending on whether the difference between (i) the mode of the read length distribution of the normal samples and (ii) the mode of the read length distribution of the tumor sample or the second most frequent read length from this distribution (if supported by at least 20% of the reads) was a multiple of three.

### Mutation calling

We utilized MuTect 1.1.4^50^ to call somatic mutations in both the tumor and matched normal whole-genome samples, using the Catalogue of Somatic Mutations in Cancer (COSMIC) v68 and dbSNP135 as reference sets of known somatic and germline mutations, respectively. To ensure the somatic origin of the variant sets reported by MuTect, we filtered out germline mutations from the 1000 Genomes Project (phase 3, release 20130502)^51^ and any mutation present in at least one read in two unmatched normal BAM files from the same tissue. Somatic mutations for all 7,919 exomes were downloaded from the GDAC (https://gdac.broadinstitute.org) website. We utilized HaplotypeCaller 3.4-46-gbc02625^52^ to examine germline mutations. We only kept deleterious mutations (i.e., frameshift, nonsense, missense and splicing site) supported by at least 10 reads, and those with at least 30% of the reads mapped to that locus supporting the alternative allele. Additionally, we only kept missense mutations with a predicted MetaLR score from Annovar^53^ higher than 0.9. We did not consider mutations in the exons 9, and 11 to 15 of *PMS2*, as the *PMS2CL* pseudogene displays more than 98% sequence identity with these exons. Due to the high allelic diversity of *PMS2CL* due to sequence transfer^24^, it is challenging to dismiss false positive mutations called in these exons.

### Correlation between gene expression and genomic events in MMR genes and proofreading polymerases

To investigate the association of the level of gene expression and genomic events on seven MMR genes (*MLH1, MLH3, MSH2, MSH3, MSH6, PMS1* and *PMS2*) and two proofreading DNA polymerases (*POLD1* and *POLE*), we utilized gene expression, promoter methylation and DNA copy number profiles for the186 MSI-H cases with available from these three data types. Gene expression profiles were first log transformed, i.e.; log_2_ (FPKM+1). Subsequently, the expression values of each row and column were median-centered and rescaled so that the sum of the squares of the values to be 1.0. To process the promoter methylation data, we collected 17 common probes corresponding to the 9 genes studied between two microarray platforms (humanmethylation27 and humanmethylation450, Illumina). β values were obtained for 17 probes and averaged per gene. The *MLH1* promoter was considered methylated in samples with β values > 0.3. To obtain copy number data, we selected segmentation files filtered for germline alterations. Log2 copy numbers overlapping the genomic segments of 8 genes were considered as the copy numbers of these genes. *POLE* was ignored since it was not covered by the segmentation files. Pearson’s correlation was used to assess the relationship between gene expression and promoter methylation (β values), as well as the relationship between gene expression and DNA copy numbers. The relationship between gene expression and somatic mutations and MSI events, was evaluated using the Mann-Whitney test (α = 0.05).

### Analysis of epigenomic features

We downloaded the coordinates of the 25-state chromatin state map defined using 12 marks (H3K4me1, H3K4me2, H3K4me3, H3K9ac, H3K27ac, H4K20me1, H3K79me2, H3K36me3, H3K9me3, H3K27me3, H2A.Z and DNase) across 127 reference epigenomes from the Epigenome Roadmap project^54^. For each of the 30 whole genomes with the highest MSI counts, the list of their MSI loci was intersected with the chromatin state maps defined using cell lines from the same anatomical location as the tumor types. We used the chromatin state maps defined using the epigenomes E092, E094, E0110 and E0111 for STAD, E117 for UCEC, E076, E106 and E075 for COAD, E086 for KICH, E027, E028 and E199 for BRCA, E053, E054, E067, E068, E069, E070, E071, E072, E073, E074, E081, E082 and E0125 for GBM, E097 for OV, E088, E096, E114 and E128 for LUSC, E055, E056, E057, E059, E061, E126, E127 and E058 for HNSC and E086 for KIRP. Subsequently, the percentage of MSI events overlapping each chromatin state was averaged across the matched epigenomes. The same process was applied to the set of MS loci from the genome-wide reference set. Fisher’s exact test was used to assess the significance of the enrichment for MSI events of each chromatin state in each of the cancer genomes. The significance level was set to 0.05.

### MSI status prediction

We used Random Forest models^40^ to build binary classifiers for the prediction of MSI status. Each tumor was encoded with a vector recording the number of MSI events and the presence or absence of MSI events in 7,863 genes targeted by MSI in at least one sample. Features displaying a variance close to zero across all learning examples (i.e., near-zero variance descriptors) were removed using the function *nearZeroVar* from the R package *caret*^55^. The remaining descriptors were mean-centered to zero and scaled to unit variance to obtain *z*-scores using the function *PreProcess* from the same package.

The final set of covariates, the MSI status of the training data and the code used to train the models are given in the Supplementary File 1-2. The number of trees was set to 100^41^, the optimal value of the parameter *mtry* was determined to be 182 *via* 10-fold cross-validation and the default values were used for the remaining parameters. With this *mtry* value, the final prediction models were built using all available learning examples.

To estimate prediction errors, we used the following pipeline^41^ from the R package *conformal* (https://cran.r-project.org/web/packages/conformal/index.html). We used cross-validation predictions to define a Mondrian class list for each category (i.e., MSI-H and MSS) by sorting in increasing order the fraction of trees voting for that class for each training example. Next, we applied the model trained on all learning examples to each sample without MSI categorization, and calculated for each case the fraction of trees in the forest voting for each class. These values were intersected with the corresponding Mondrian class list. For each sample, the *P* value for a given class was calculated as the number of elements in the corresponding Mondrian class list higher than the vote fraction for that class divided by the number of elements in that list. If the *P* value for a given class is above the tolerated error, ε, the sample is predicted to belong to that category. Hence, a given sample may be called as MSI-H or MSS. However, it can also be called as both in cases when the model does not have enough predictive power to discriminate between classes, or neither in cases when the sample is outside the applicability domain of the model. This flexibility thus gives an unbiased estimate of the reliability of the predictions given the training data. The tolerated error, ε, indicates the maximum fraction of predictions that are incorrect. Therefore, increasing the confidence level might increase the number of uncertain predictions, that is, samples classified as both MSI-H and MSS.

## Acknowledgements

The results published here are based upon data generated by The Cancer Genome Atlas and obtained from the Database of Genotypes and Phenotypes (dbGaP) with accession number phs000178.v8.p7. Information about TCGA can be found at http://cancergenome.nih.gov. We thank the Research Information Technology Group at Harvard Medical School and KT for providing computational resources. This work was supported by grants from the Ludwig Center at Harvard (I.C.C. and P.J.P.) and the Korean Health Technology R&D Project, Ministry of Health & Welfare, Republic of Korea (HI13C2096, S.L. and W.-Y.P.). We thank Alison Barton and Lixing Yang for their critical reading of the paper.

## Author contributions

S.L. and I.C.C. performed bioinformatic analysis of all data, with guidance from T.M. and P.J.P. S.L. was supervised by W.-Y.P.; I.C.C. was supervised by P.J.P. The manuscript was written by I.C.C. and P.J.P. with substantial input from T.K. and S.L.

## Supplementary Figures

**Figure S1.**
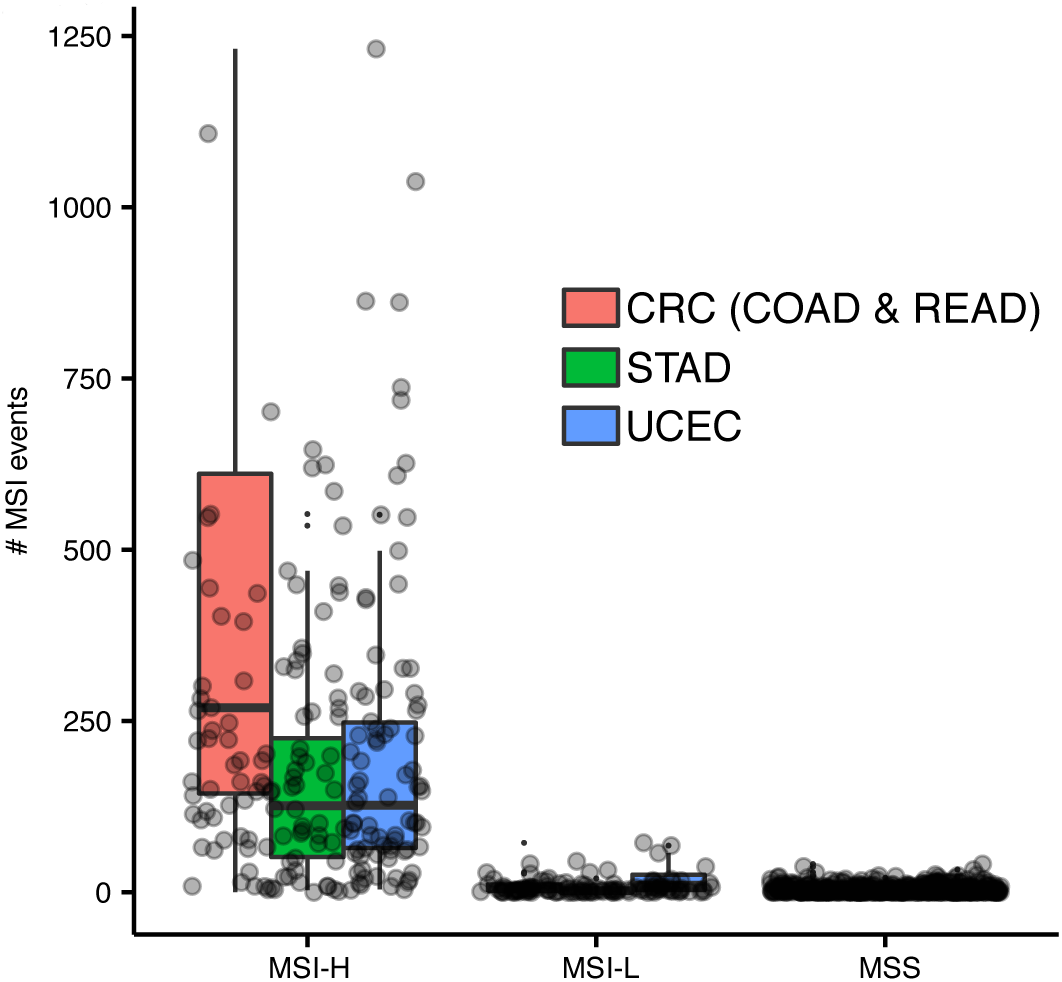
Distribution of MSI events in MSI-H, MSI-L and MSS cases in MSI-prone tumor types.

**Figure S2.**
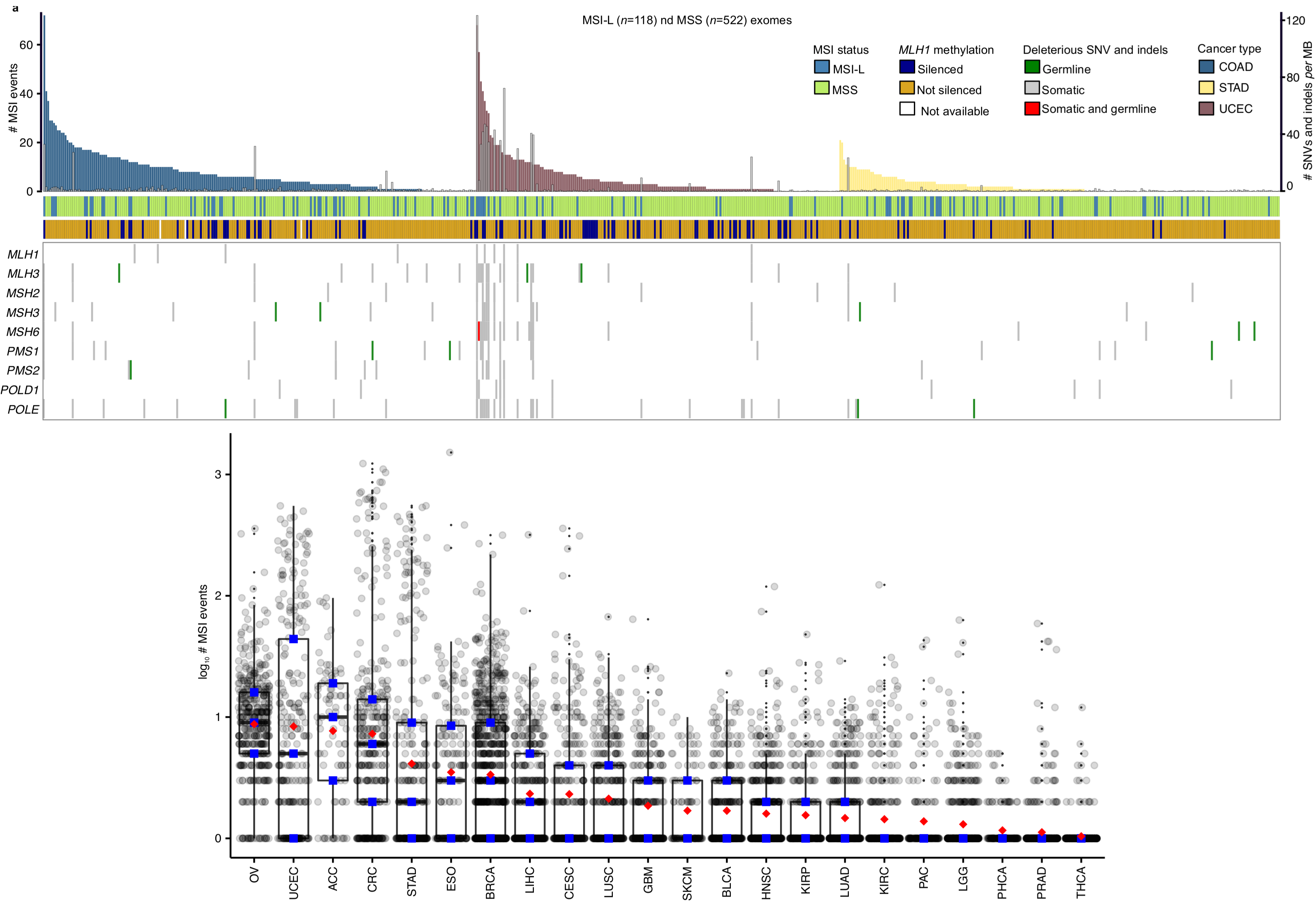
(**a**) Landscape of MSI in MSI-L and MSS tumors. (**b**) Distribution of MSI events in low-frequency MSI-H tumor types. Blue points indicate the median and the interquartile range (25^th^ through 75^th^ percentile), whereas red points indicate the mean value.

**Figure S3.**
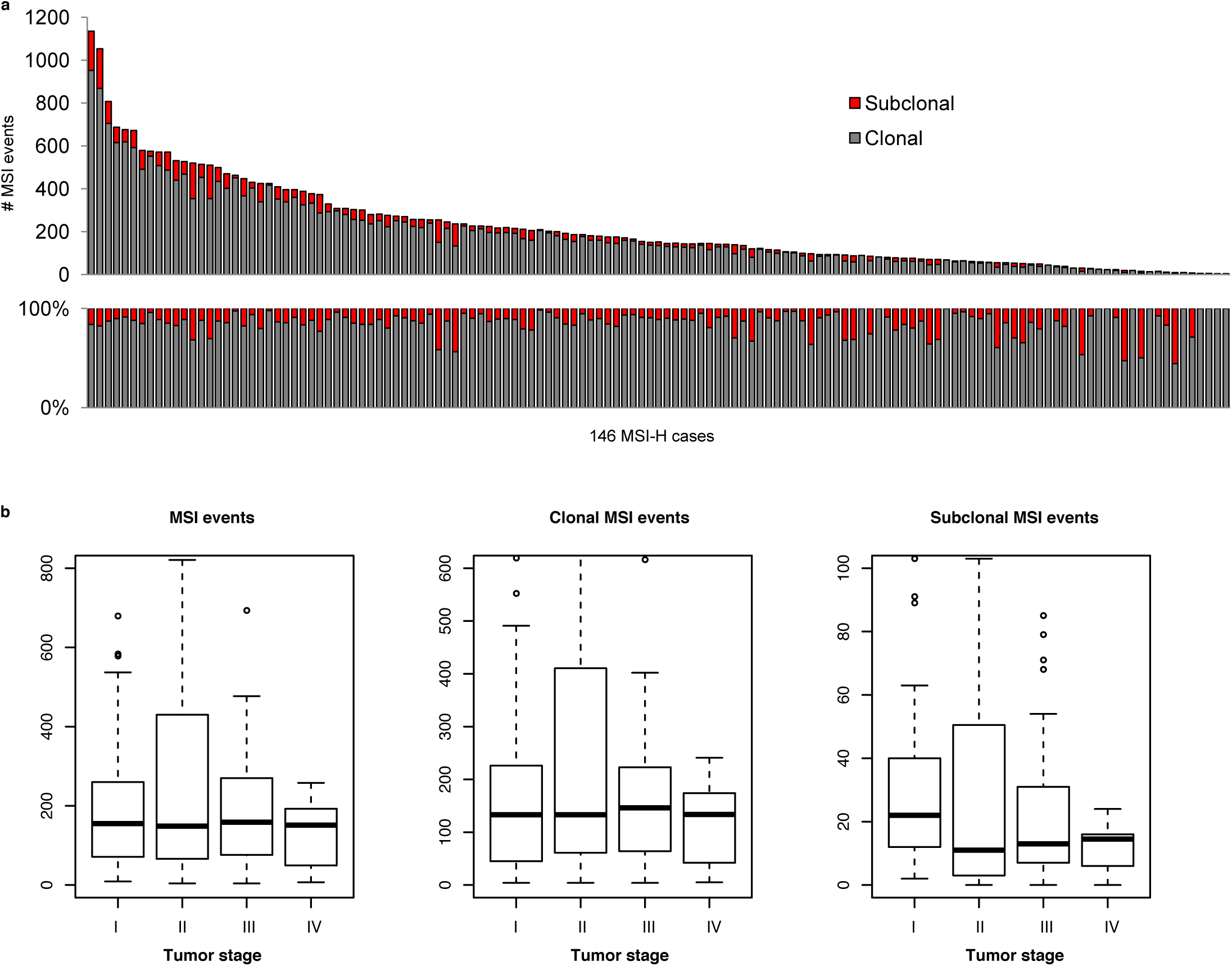
The abundance of clonal and subclonal MSI events is shown for 146 MSI-H cases (**a**). Distribution of clonal and subclonal MSI events across tumor stages.

**Figure S4.**
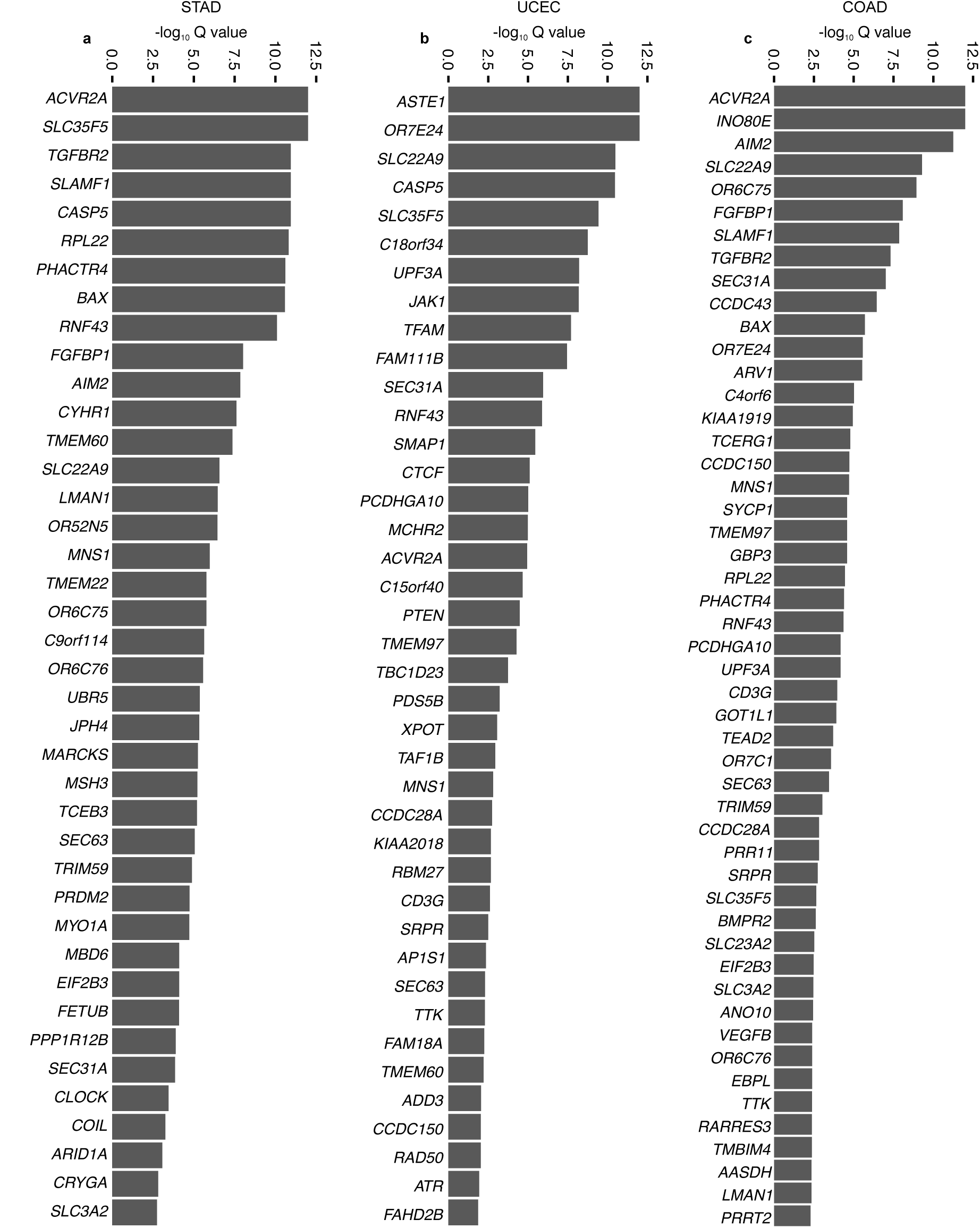
Top-ranked genes with significantly recurrent MSI in STAD (**a**), UCEC (**b**) and COAD (**c**) tumors (Q value, FDR < 0.05; MutSigCV).

**Figure S5.**
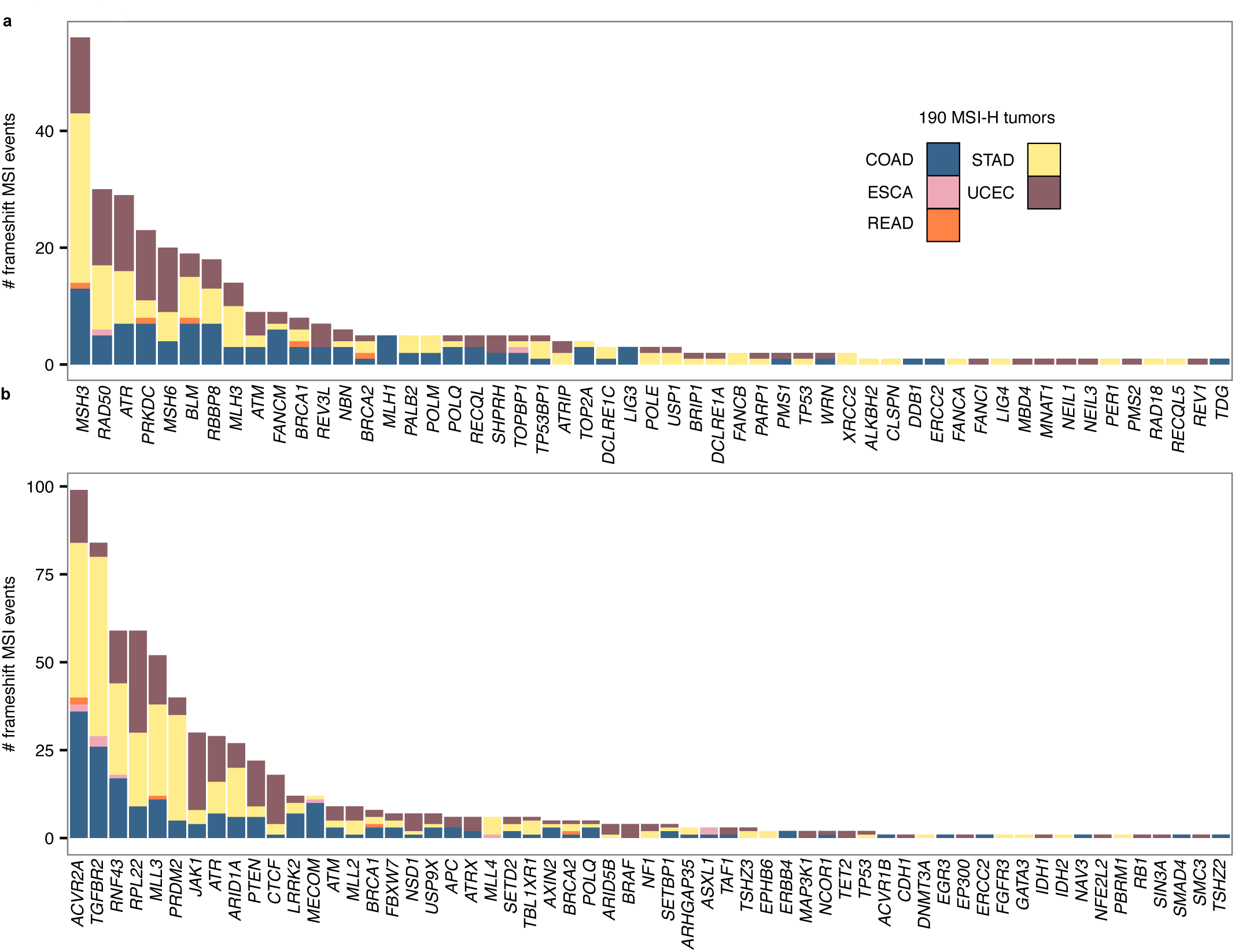
Abundance of frameshift MSI events in genes implicated in DNA repair (**a**) and tumorigenesis (**b**) across 5 MSI-prone cancer types.

**Figure S6.**
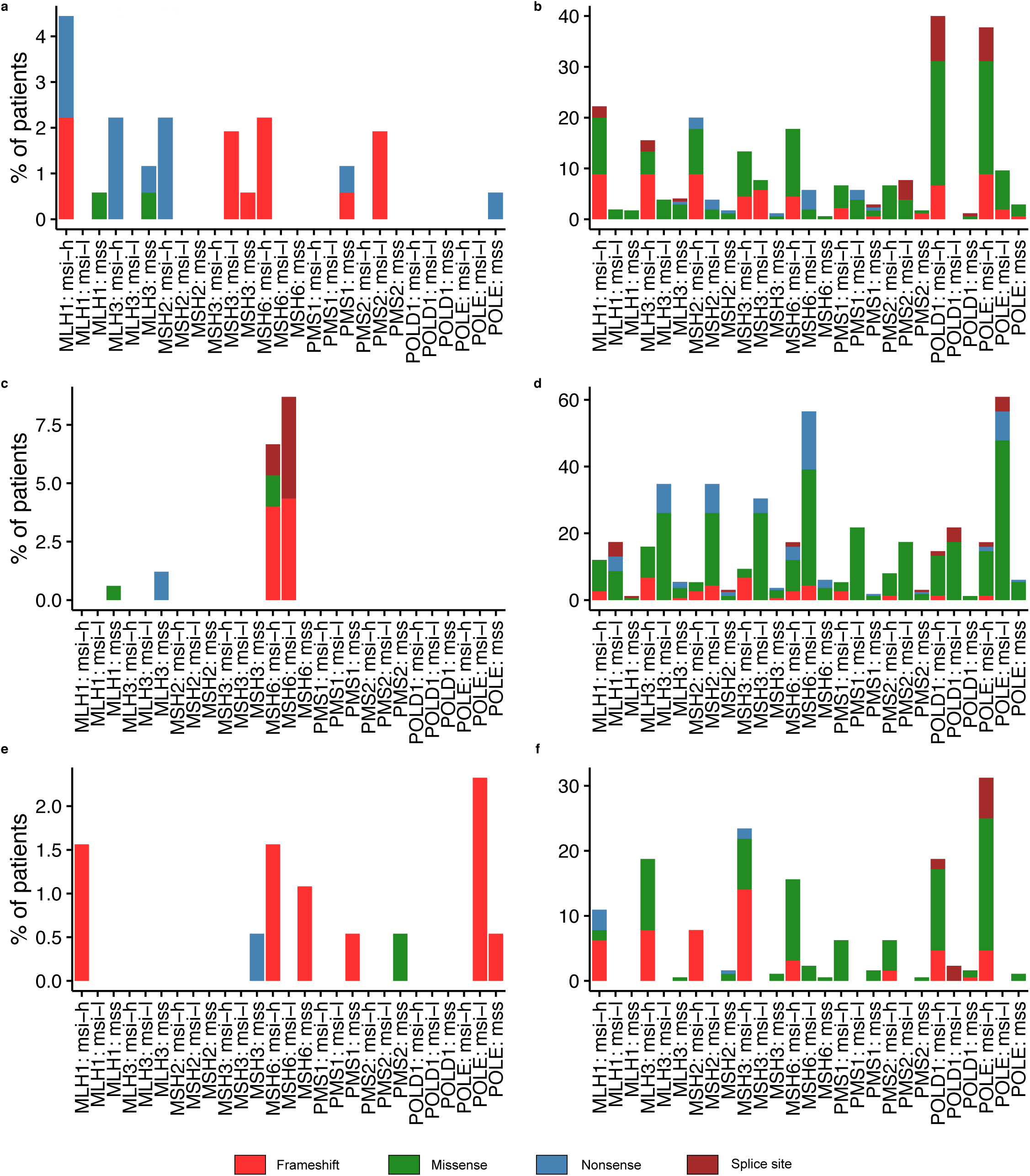
Frequency of SNVs and indels in MMR genes, *POLE* and *POLD1* in COAD (**a,b**), UCEC (**c,d**) and STAD (**e,f**).

**Figure S7.**
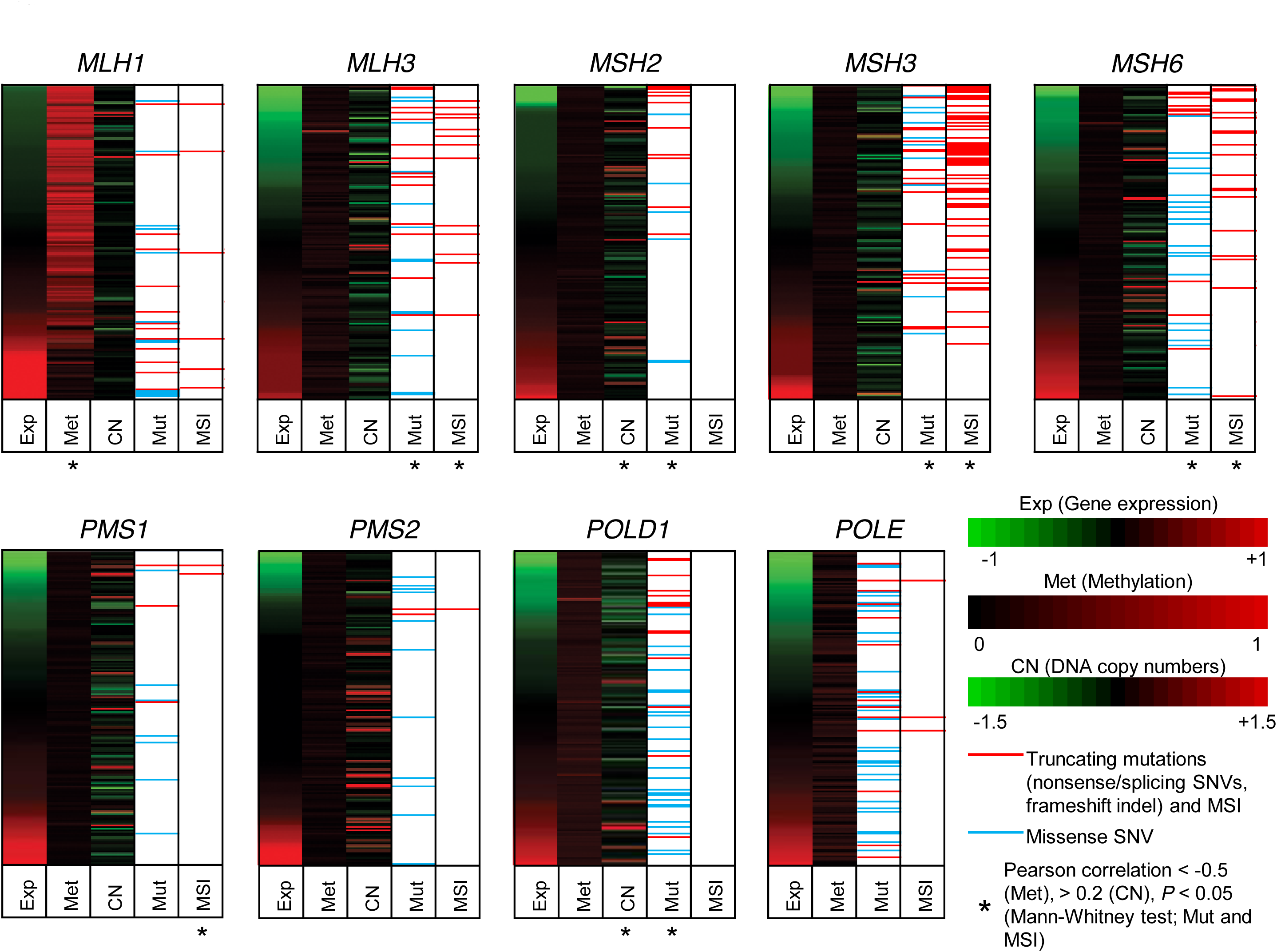
Gene expression, promoter methylation and DNA copy number profiles for MMR genes (*MLH1, MLH3, MSH2, MSH3, MSH6, PMS1* and *PMS2*) and two proofreading DNA polymerase (*POLD1* and *POLE*) corresponding to186 MSI-H cases.

**Figure S8.**
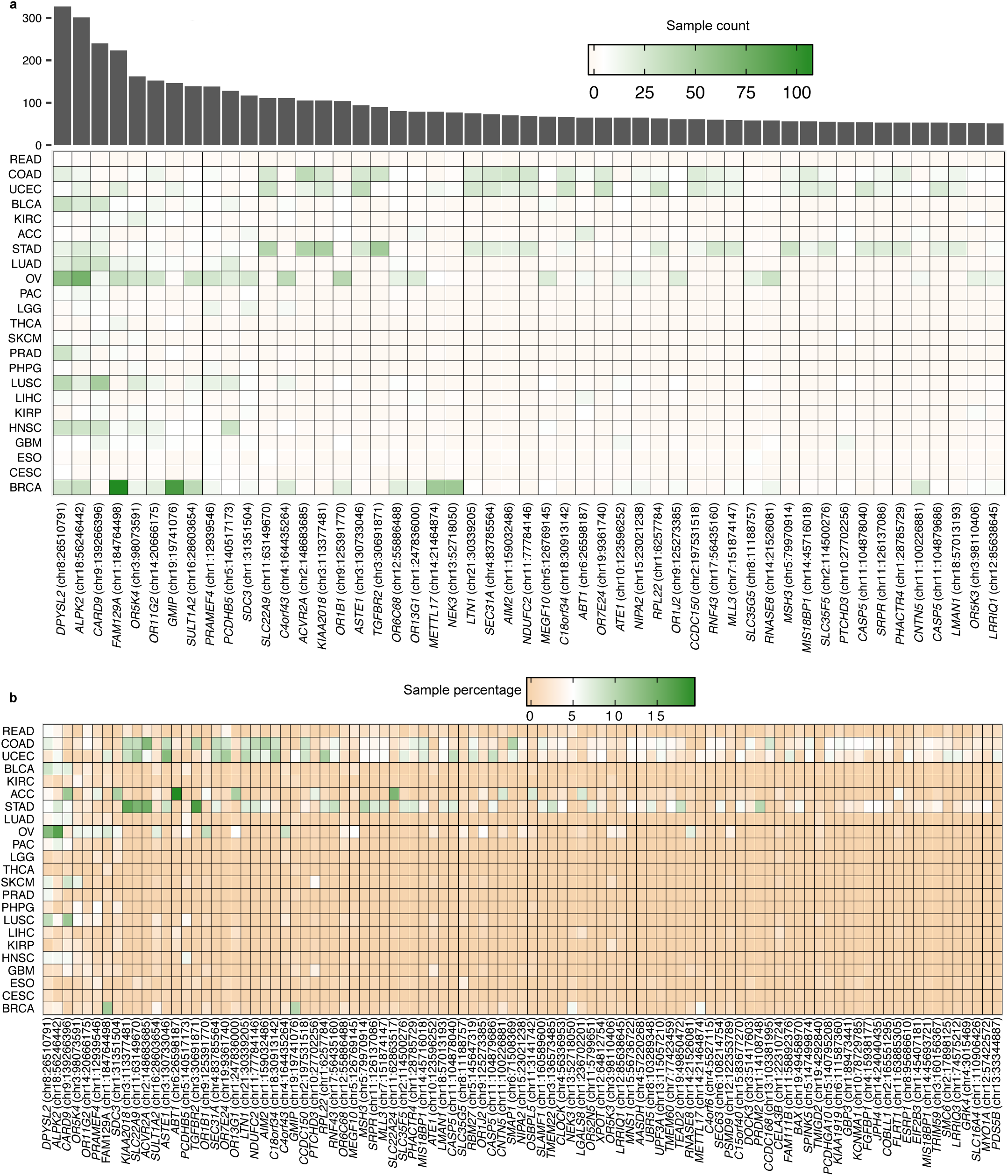
Coding MSI loci recurrently subject to frameshift MSI events in the 22 cancer types studied (Table 1). This analysis included MSI-H, MSI-L and MSS tumors. The heatmap shows the fraction of tumors from each cancer type harboring frameshift MSI mutations in MS loci located within the coding sequence of the genes indicated on the *x* axis. The total count of frameshift MSI events at those MSI loci across all tumor types is depicted in the above barplot.

**Figure S9.**
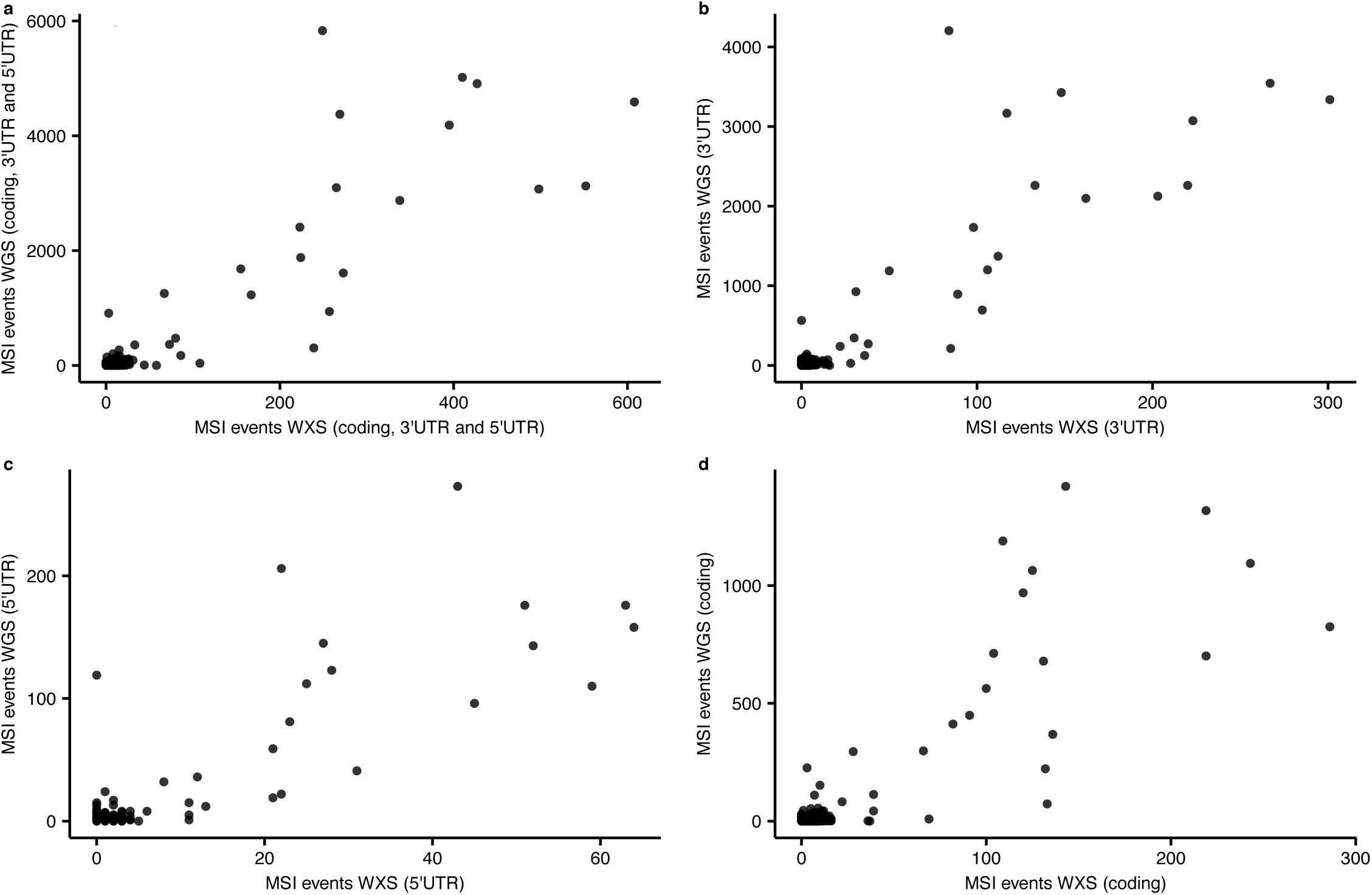
Cross-platform comparison of MSI calls in exonic (coding and 3’/5’ UTR) (**a**), 3’UTR (**b**), 5’UTR (**c)** and coding (**d**) MS repeats.

**Figure S10.**
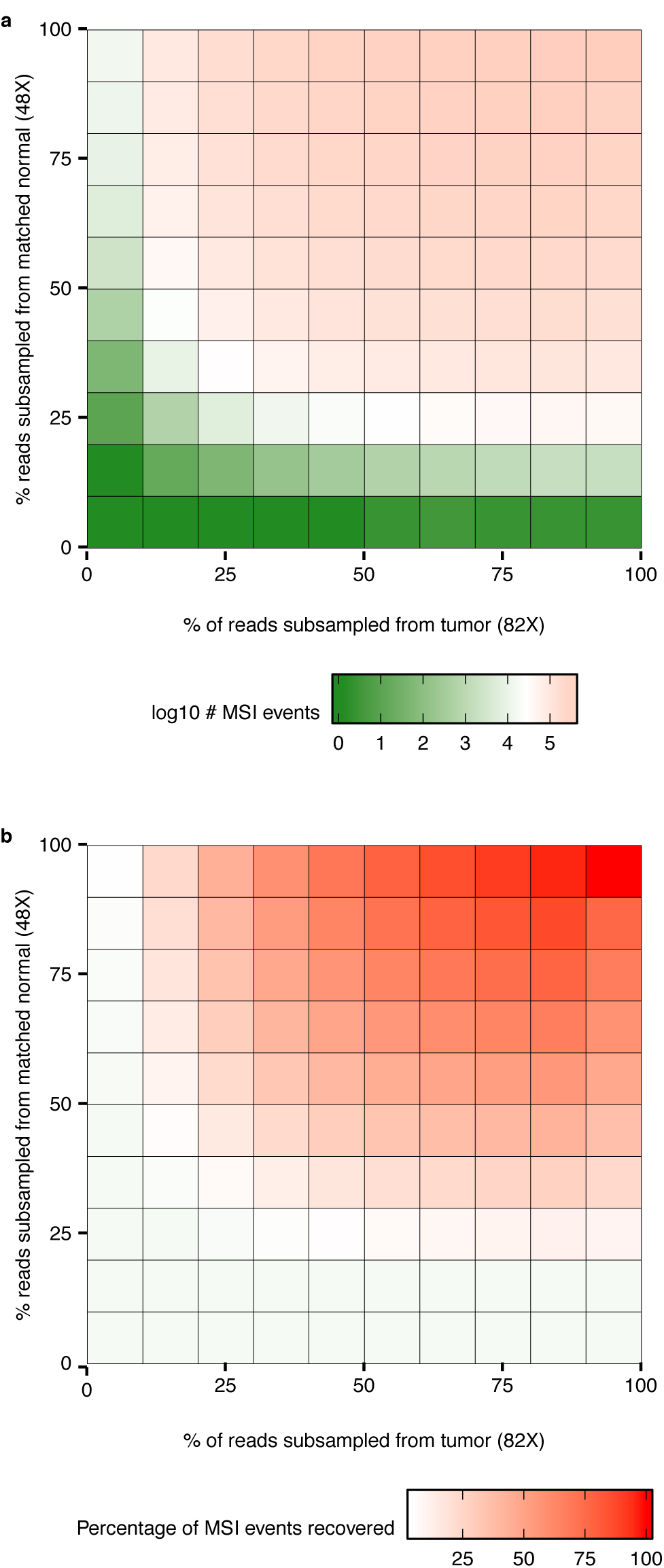
Influence of read depth on MSI calling across increasing larger random subsamples of reads from the tumor (82X) and matched normal (42X) bam files corresponding to the patient TCGA-AD-A5EJ. The total number of MSI events and the percentage of MSI called at different subsampling levels are depicted in (**a**) and (**b**), respectively.

**Figure S11.**
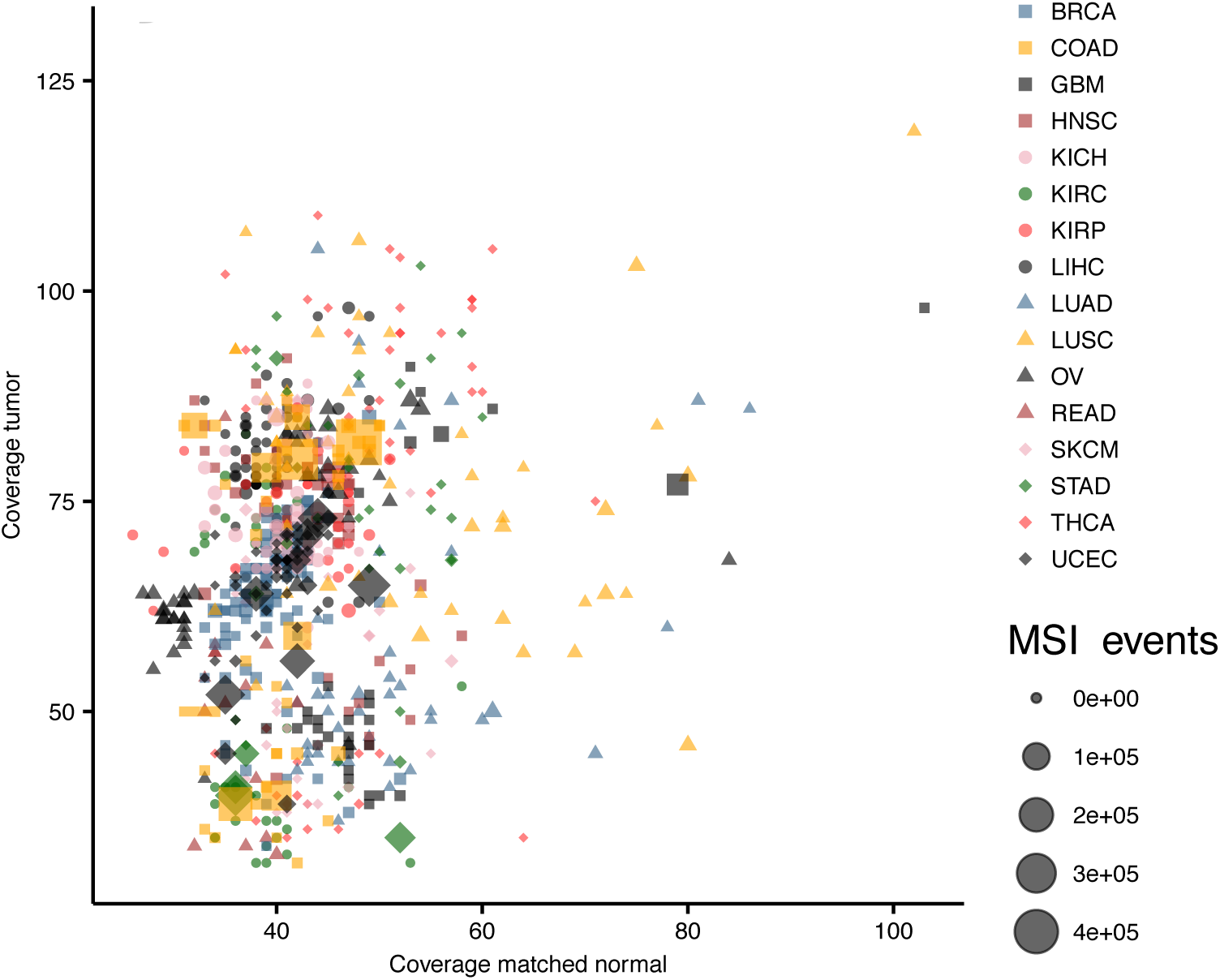
Average read depth of the tumor (y-axis) and matched normal (x-axis) samples.

**Figure S12.**
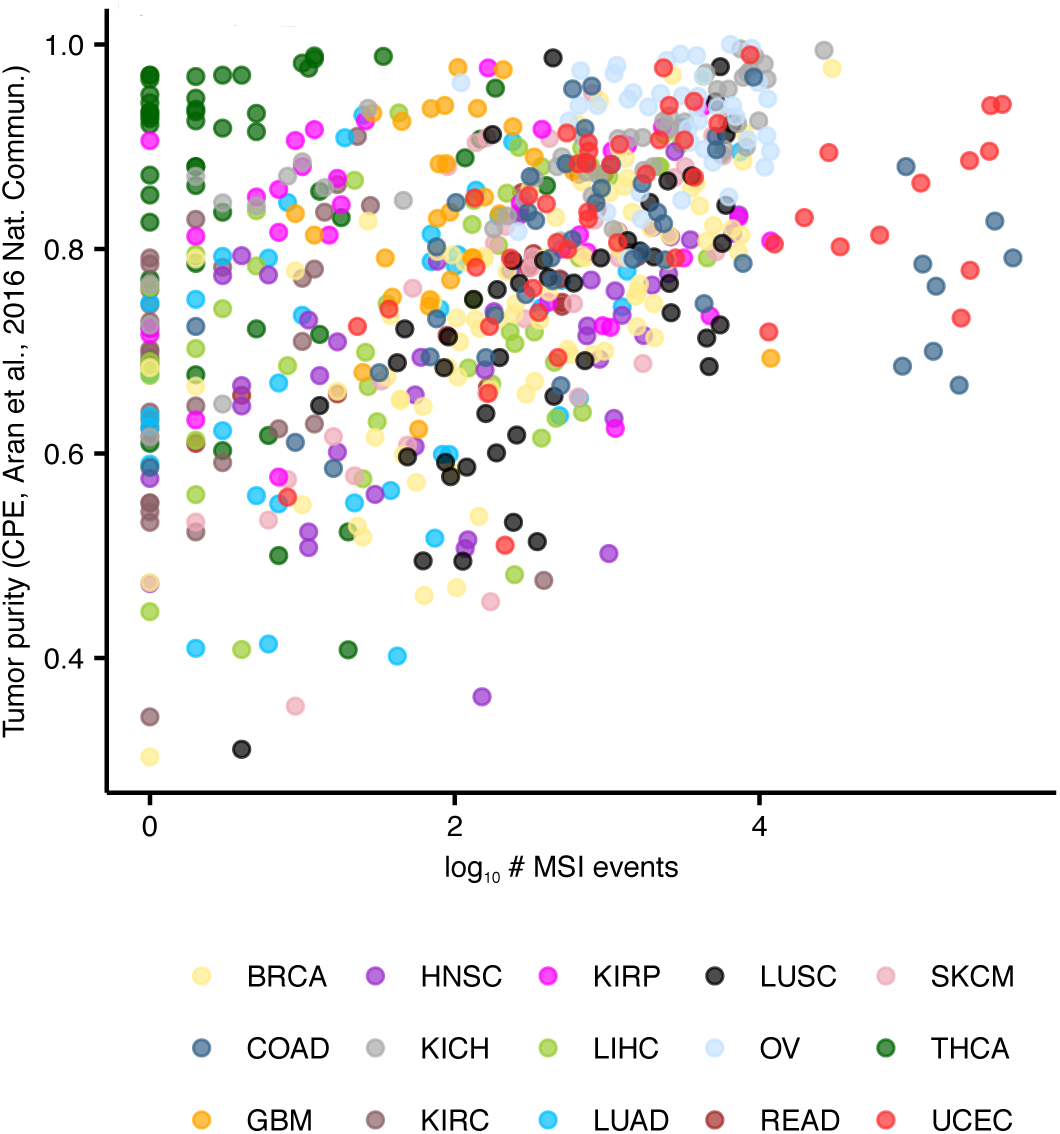
Correlation between MSI rates *vs* tumor purity.

**Figure S13.**
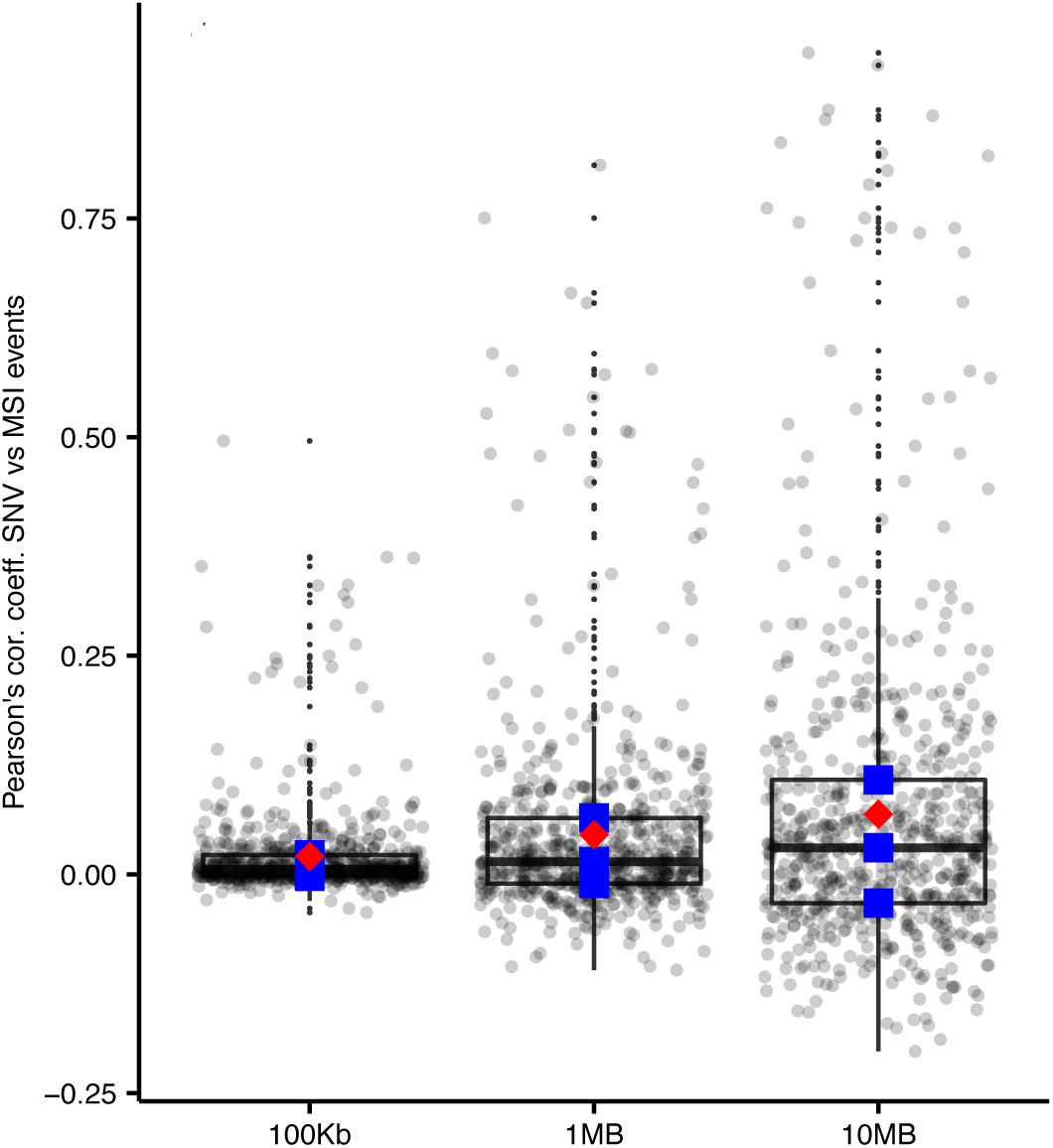
Distribution of Pearson correlation coefficients for SNV and MSI rates measured in (**a**) 1Mb, (**b**) 10Mb and (**c**) 100Kb bins. Blue points indicate the median and the interquartile range (25th-75th percentile), whereas red points indicate the mean value.

**Figure S14.**
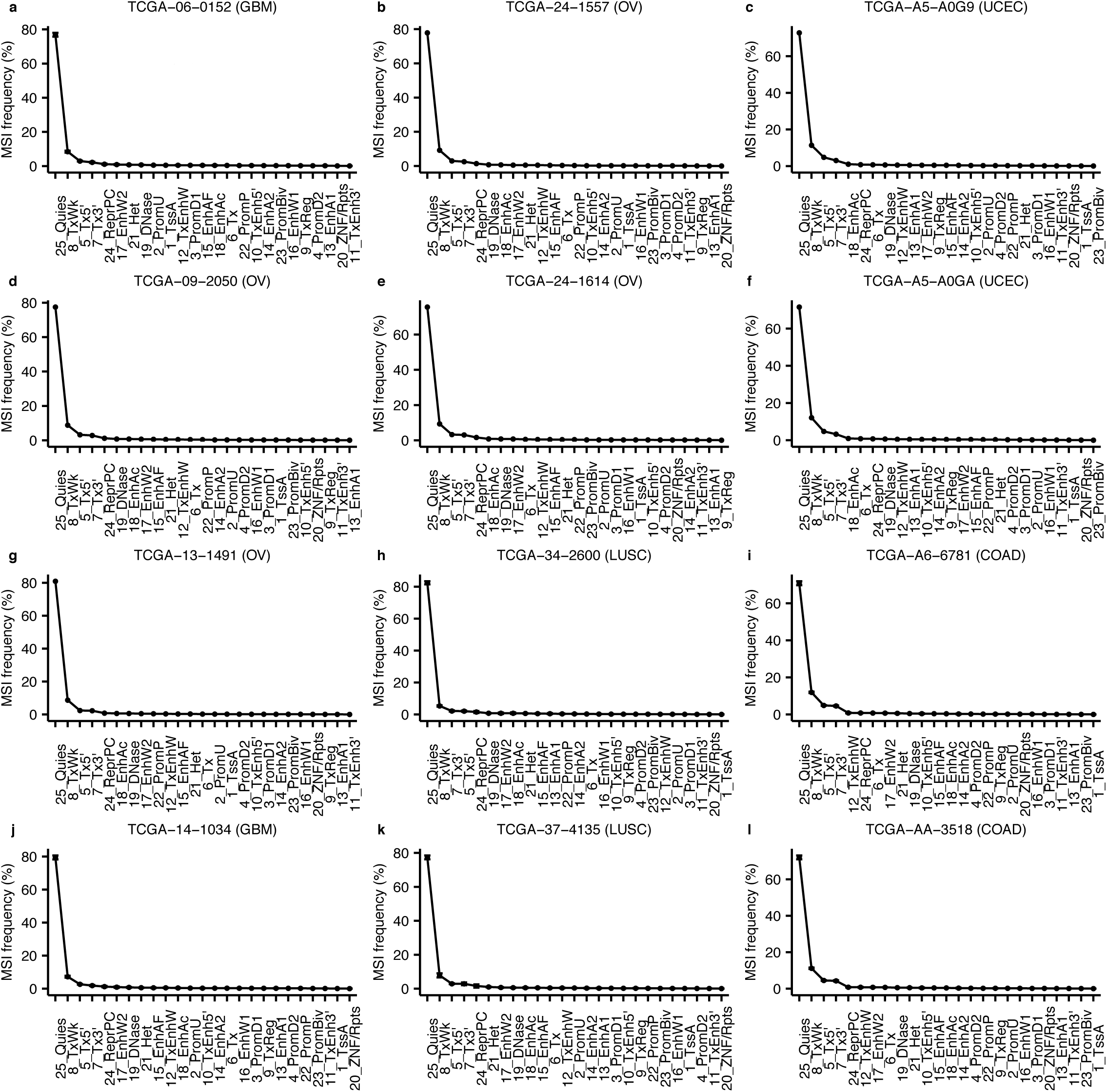
Frequency of MSI events in the 25 states of the chromatin state map averaged across 127 epigenomes for samples TCGA-06-0152 (**a**), TCGA-24-1557 (**b**), TCGA-A5-A0G9 (**c**), TCGA-09-2050 (**d**), TCGA-24-1614 (**e**), TCGA-A5-A0GA (**f**), TCGA-13-1491 (**g**), TCGA-34-2600 (**h**), TCGA-A6-6781 (**i**), TCGA-14-1034 (**j**), TCGA-37-4135 (**k**) and TCGA-AA-3518 (**l**).

**Figure S15.**
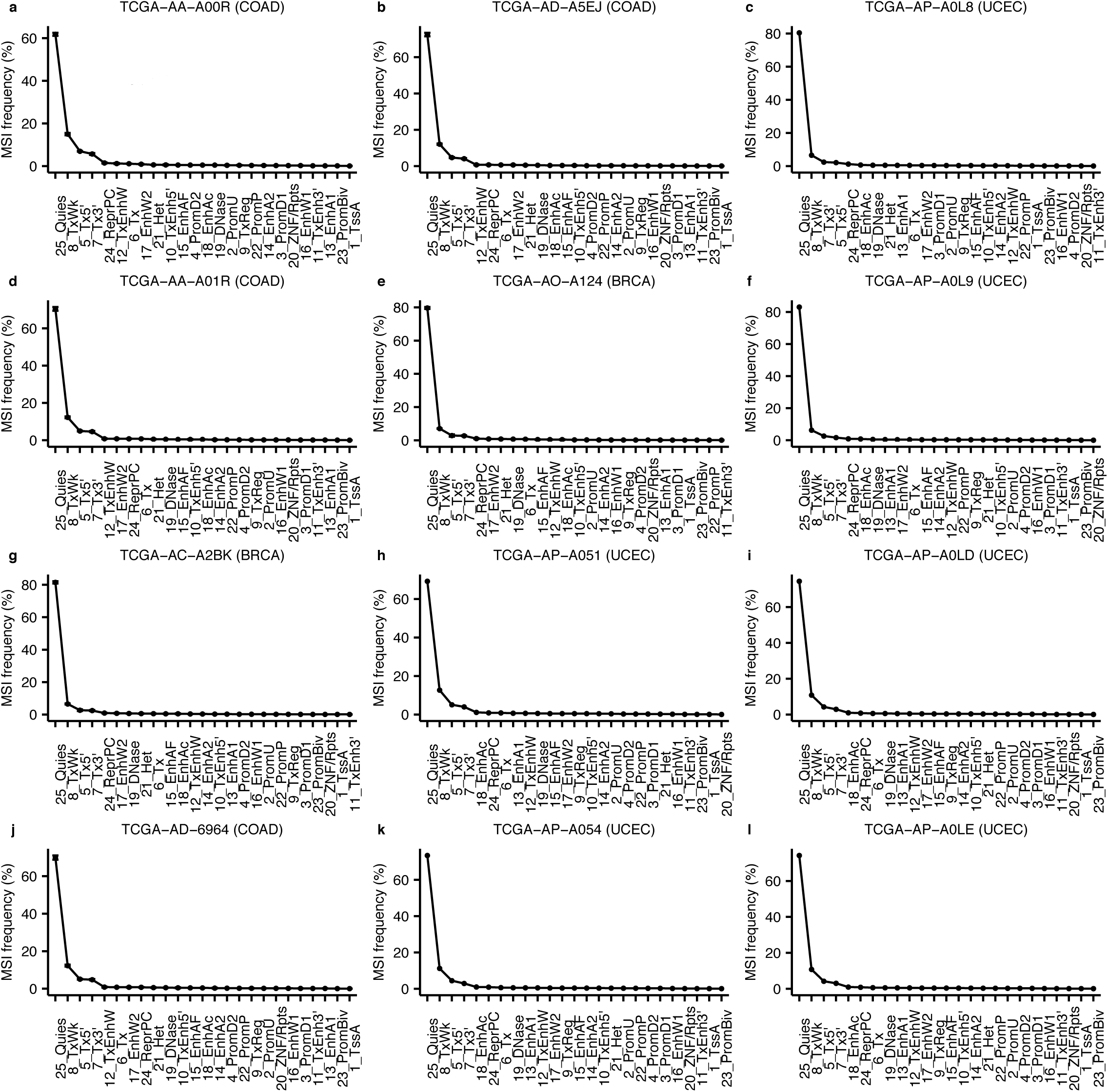
Frequency of MSI events in the 25 states of the chromatin state map averaged across 127 epigenomes for samples TCGA-AA-A00R (**a**), TCGA-AD-A5EJ (**b**), TCGA-AP-A0L8 (**c**), TCGA-AA-A01R (**d**), TCGA-AO-A124 (**e**), TCGA-AP-A0L9 (**f**), TCGA-AC-A2BK (**g**), TCGA-AP-A051 (**h**), TCGA-AP-A0LD (**i**), TCGA-AD-6964 (**j**), TCGA-AP-A054 (**k**) and TCGA-AP-A0LE (**l**).

**Figure S16.**
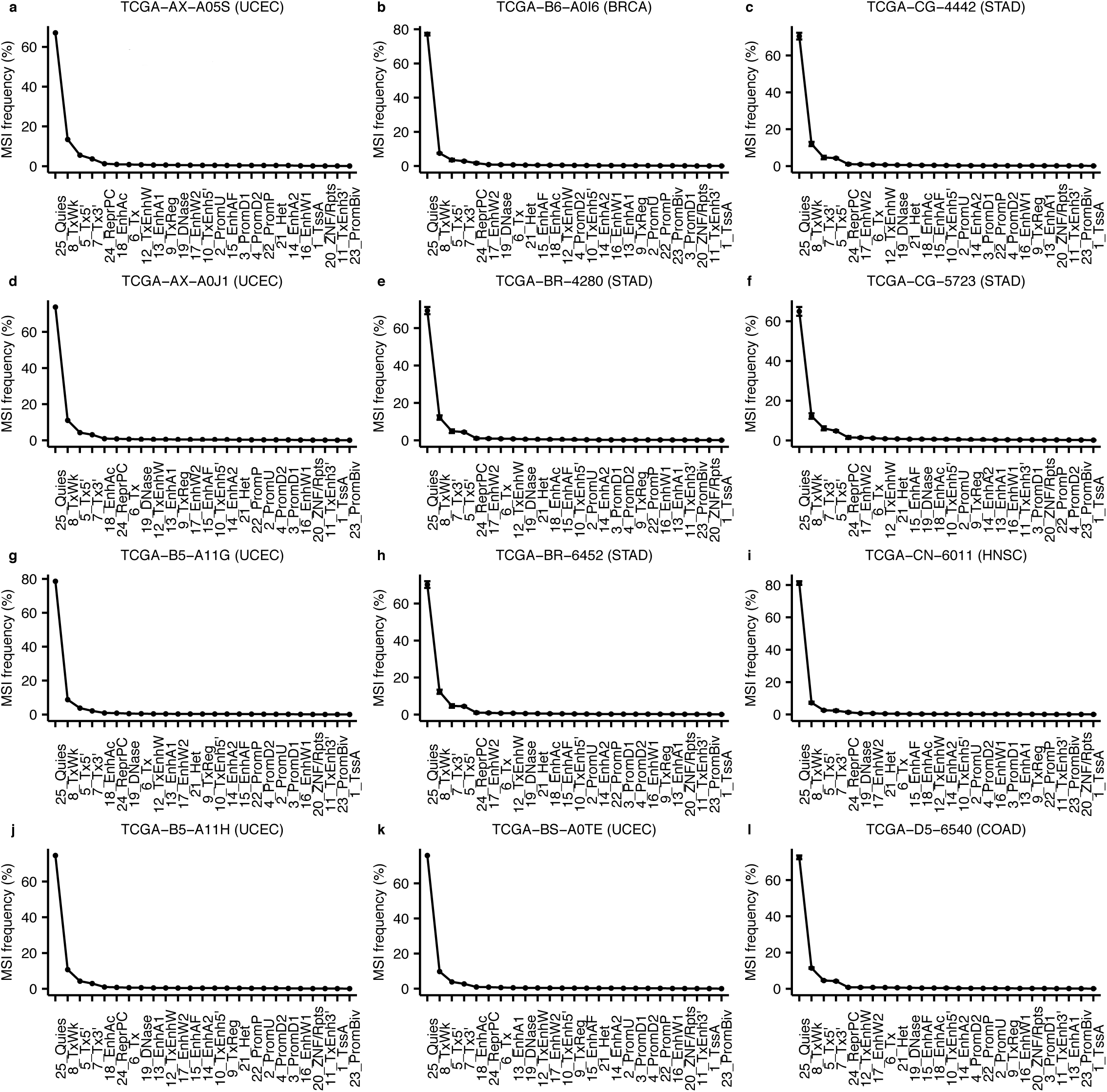
Frequency of MSI events in the 25 states of the chromatin state map averaged across 127 epigenomes for samples TCGA-AX-A05S (**a**), TCGA-B6-A0I6 (**b**), TCGA-CG-4442 (**c**), TCGA-AX-A0J1 (**d**), TCGA-BR-4280 (**e**), TCGA-CG-5723 (**f**), TCGA-B5-A11G (**g**), TCGA-BR-6452 (**h**), TCGA-CN-6011 (**i**), TCGA-B5-A11H (**j**), TCGA-BS-A0TE (**k**) and TCGA-D5-6540 (**l**).

**Figure S17.**
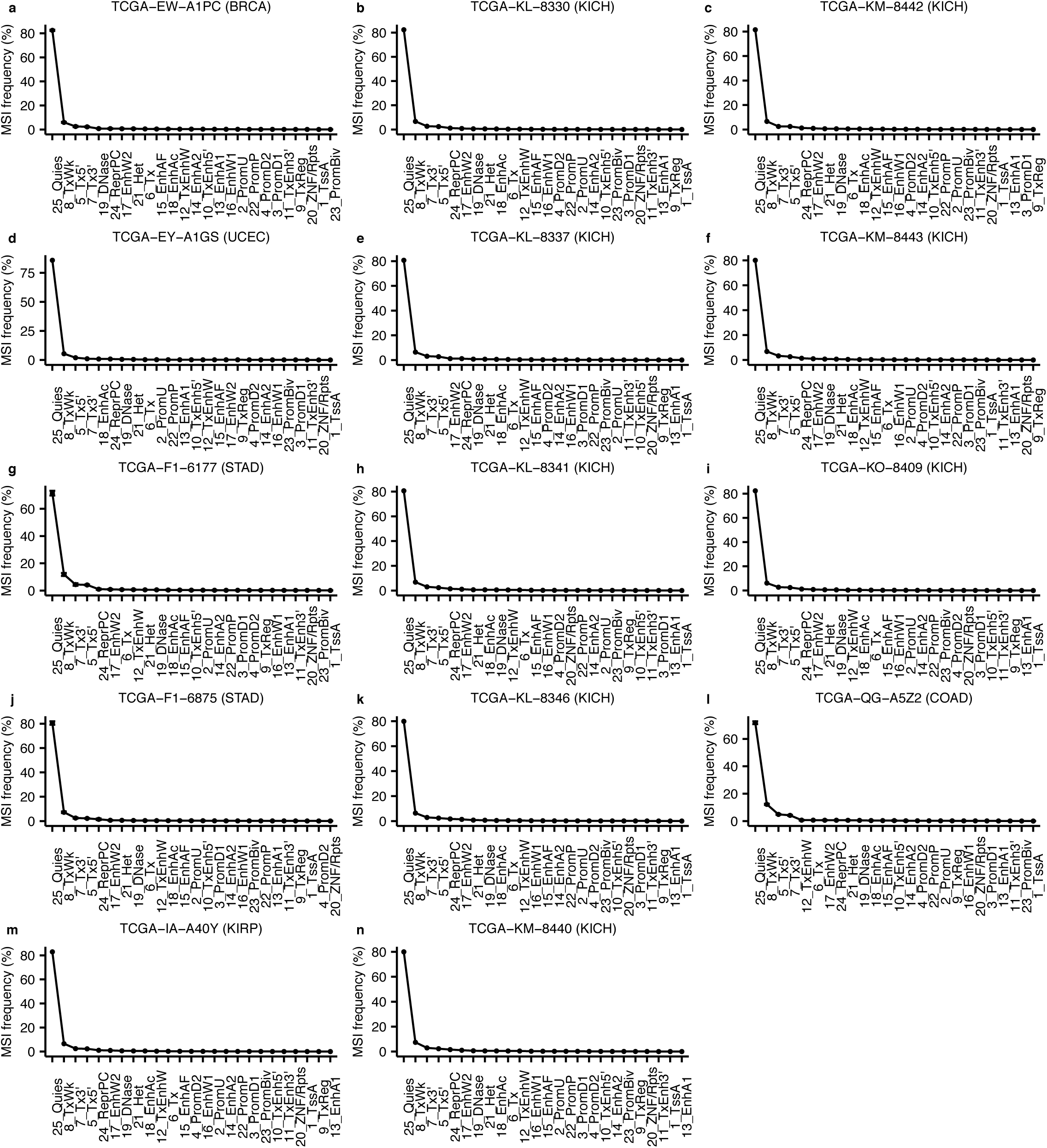
Frequency of MSI events in the 25 states of the chromatin state map averaged across 127 epigenomes for samples TCGA-EW-A1PC (a), TCGA-KL-8330 (b), TCGA-KM-8442 (c), TCGA-EY-A1GS (d), TCGA-KL-8337 (e), TCGA-KM-8443 (f), TCGA-F1-6177 (g), TCGA-KL-8341 (h), TCGA-KO-8409 (i), TCGA-F1-6875 (j), TCGA-KL-8346 (k), TCGA-QG-A5Z2 (l), (**m**) TCGA−IA−A40Y and (**n**) TCGA−KM−8440.

**Figure S18.**
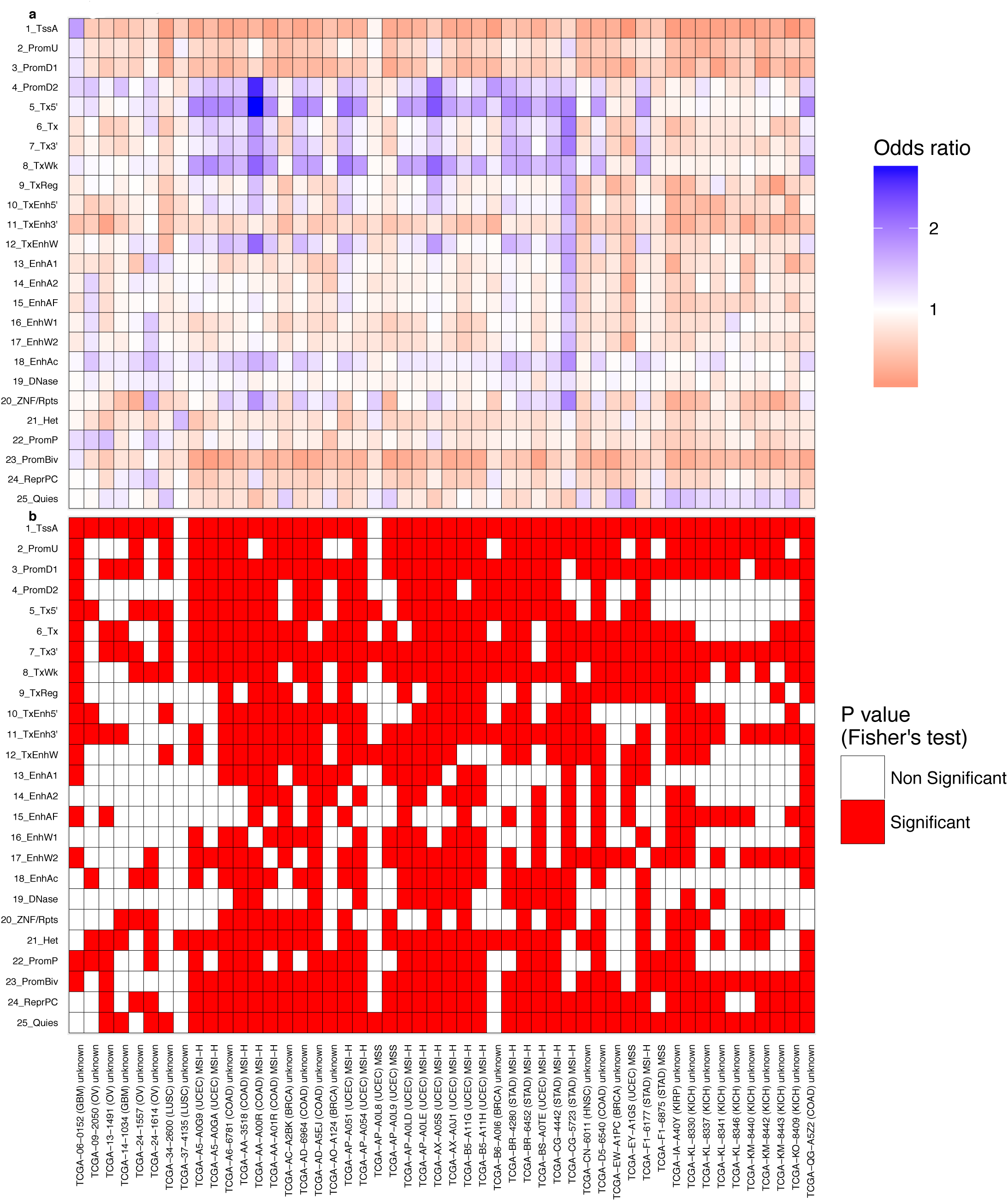
Enrichment of MSI events in the 25 states of the chromatin state map. Odds ratio (**a**) and p values (**b**) for each chromatin state in the 30 whole-genomes displaying the highest MSI rates. Enrichment of MSI events in each chromatin state was tested using Fisher’s exact test (Online Methods).

**Figure S19.**
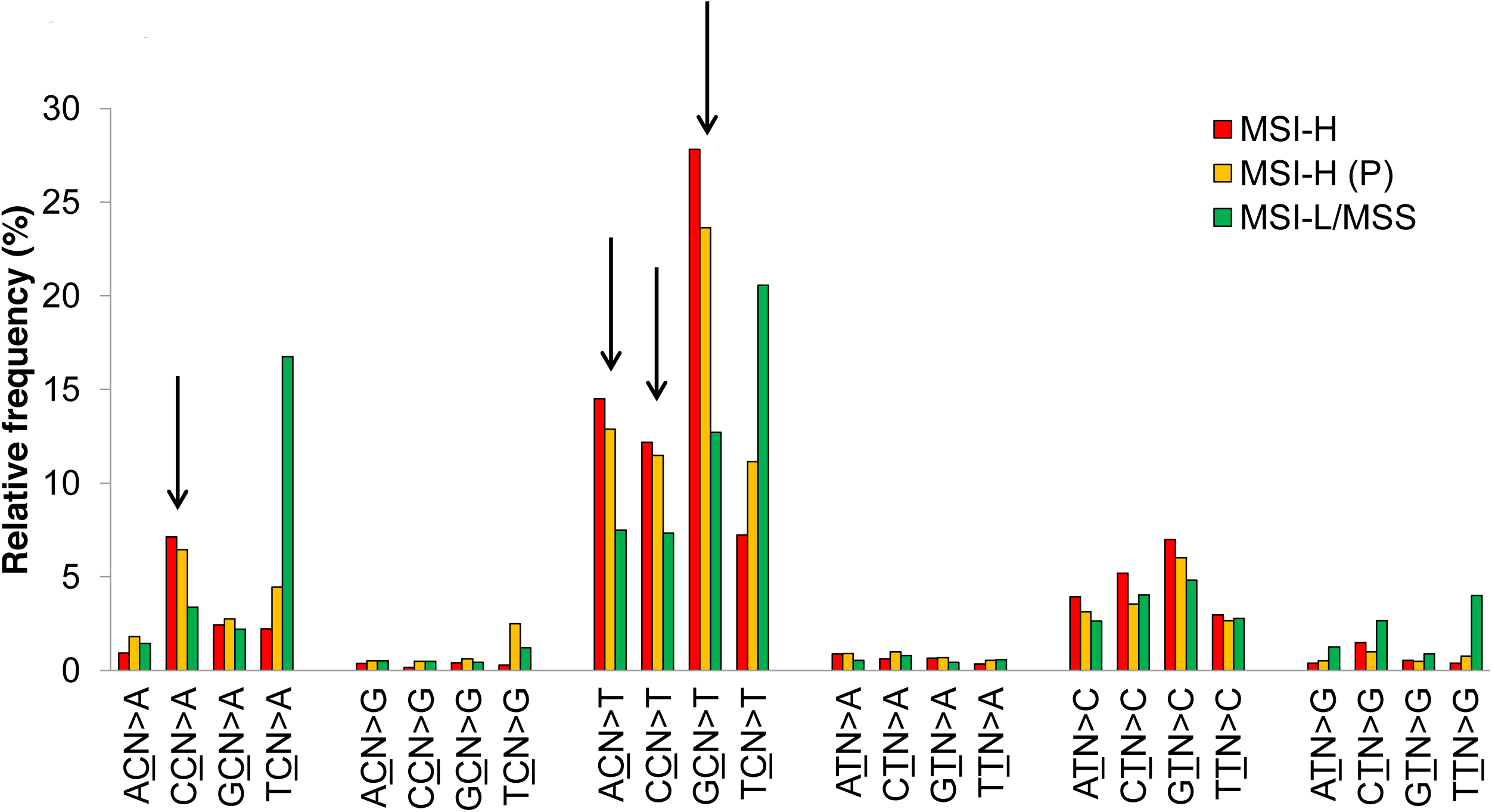
Mutation signatures of predicted MSI-H genomes. The relative frequencies of the 96 trinucleotide mutation contexts (strand symmetric) are shown for MSI-H, MSI-H (predicted) and MSI-L/MSS cases (red, orange and green, respectively). Mutations were pooled across the genomes of three MSI-prone tumor types of COAD/READ, STAD and UCEC. Arrows indicate known mutation features of MSI-H genomes, e.g., C>T transitions in (A/C/G)pCpN sequence contexts and C>A transversions at an CpCpN context, which are enriched in both MSI-H and MSI-H (predicted) genomes compared to MSI-L/MSS genomes.

## Supplementary Tables

**Table S1** The TCGA barcodes and the total number of MSI calls showing significant differences (FDR < 0.05; Kolmogorov-Smirnov test) in the MS allele lengths between the tumor and matched normal exomes are listed.

**Table S2** Results of the DAVID analysis.

**Table S3** Frameshift MSI events observed in MSI-H tumors among 13 out the 151 DNA repair genes examined are listed.

**Table S4** Frameshift MSI events observed in MSI-H tumors among 12 out of the 130 major cancer-related genes examined are listed.

**Table S5** The coding MS loci harboring recurrent frameshift MSI events in COAD, STAD and UCEC MSI-H tumors found using exome sequencing data are listed.

**Table S6** Recurrent MSI events in 3’-UTR regions are listed.

**Table S7** Recurrent MSI events in 5’-UTR regions are listed.

**Table S8** The coding MS repeats harboring recurrent frameshift MSI events in MSS, MSI-L and MSI-H genomes identified using whole exome sequence data from 22 cancer types are listed.

**Table S9** Enrichment for frameshit, 3’ UTR and 5’ UTR MSI events in COAD, UCEC and STAD. The significance was calculated using the one-tailed exact Fisher’s text (α = 0.05).

**Table S10** The barcodes, the total number of significant MSI calls (FDR < 0.05; Kolmogorov-Smirnov test), and the total number of MS loci profiled are listed for the 708 whole-genomes analyzed.

**Table S11** List of MS loci discovered in the human mitochondrial DNA.

**Table S12** The barcodes corresponding to the samples used in the analysis of MSI in mitochondrial DNA are listed.

**Table S13** Top 1,000 most recurrently altered MS loci in COAD, STAD and UCEC MSI-H exomes.

**Table S14** Top 1,000 most recurrently altered MS loci in COAD, STAD and UCEC MSI-H whole genomes.

**Table S15** Predicted MSI status for 7,096 exomes.

**Table S16** The exonic MS loci discovered in the human genome are listed.

**Table S17** Genome-wide reference set of MS repeats.

**Supplementary File S1** Matrix of covariates used to train the Random Forest models.

**Supplementary File S2** Code implemented to predict MSI status using Random Forest models and conformal prediction.

